# Untangling structural molecular details of the endocytic adaptor protein CALM upon binding with phosphatidylinositol 4,5-bisphosphate-containing model membranes

**DOI:** 10.1101/2024.12.28.630489

**Authors:** Andreas Santamaria, Daniel Pereira, Nisha Pawar, Bernard T. Kelly, Javier Carrascosa-Tejedor, Mariana Sardo, Luís Mafra, Giovanna Fragneto, David J. Owen, Ildefonso Marín-Montesinos, Eduardo Guzmán, Nathan R. Zaccai, Armando Maestro

## Abstract

Clathrin assembly lymphoid myeloid leukemia protein (CALM) is involved in the formation of clathrin-mediated endocytic coats by virtue of binding many proteins involved in the process, including clathrin itself and AP2 cargo adaptor complex. CALM is able to specifically recognize the inner leaflet of the plasma membrane by binding the membrane’s phosphatidylinositol 4,5-bisphosphate (PtdIns(4,5)P_2_). Here, a biophysical approach, primarily using neutron and X-ray scattering and solid-state NMR experiments, was exploited to investigate CALM interaction with PtdIns(4,5)P_2_-presenting model membranes. The presented experimental data reveal how the CALM folded domain is partly accommodated within the lipid membrane, directly interacting with PtdIns(4,5)P_2_ phosphates. Moreover, these data suggest that CALM’s amphiphilic N-terminal helix buries into the membrane, not only stabilising the protein docking to the membrane but also providing a mechanism to induce membrane curvature.

## Introduction

Clathrin-mediated endocytosis (CME), is the main endocytic pathway used by cells to internalize membrane proteins, commonly called cargoes^1–3^, and hence to control the plasma membrane proteome. CME depends on the interaction of various cargo/adaptor and regulatory/access proteins with the phosphatidylinositol 4,5-bisphosphate (PtdIns(4,5)P_2_)-rich inner leaflet of the plasma membrane. Adaptor proteins, such as AP2 (Adaptor protein complex 2) and CALM (clathrin assembly lymphoid myeloid leukemia), bridge clathrin and cargoes embedded in the plasma membrane. Recruitment of clathrin trimers onto clathrin adaptors pre-localised at the plasma membrane leads to their self-assembly into a polygonal lattice. This lattice maintains a high local concentration of cargo adaptors and their bound cargo as the membrane under the clathrin lattice deforms inwards to produce an invaginated clathrin-coated pit (CCP). This structure is then ‘pinched off’ the membrane to form a clathrin-coated vesicle (CCV) which is subsequently transported into the cell ^4,5^. Malfunctioning during any step of this complex biological machinery can result in neurodegenerative diseases such as Alzheimer^6^ and Stiff man syndrome^7^, among other ^8,9^.

CALM is one of the most abundant clathrin adaptors in endocytic CCVs isolated from tissue culture cells (30%-35% of the adaptors in a CCV^10,11^). It has an essential role in binding to and sorting the small proteins R-SNAREs VAMPs 2, 3, and 8 which participate in the ultimate fusion of the CCV with its target membrane ^6,12^. CALM comprises a relatively small ∼30kDa , compact stacked-helical domain, called AP180 N-terminal homology (ANTH) domain (residues 19-289 ^6,13^), and a large, intrinsically unstructured C-terminal appendage containing a clathrin binding site^13–15^. Both PtdIns(4,5)P_2_ and cargo binding sites are located in the ANTH domain, involving histidine 41, lysines 28, 38, 40^13^ and methionine 244^6^, respectively^16^. CALM is proposed to possess an amphipathic helix at the N-terminus (amphipathic helix 0, AH0, residues 5-18)^16^, which orientation is hypothesised to change upon membrane binding^16^. CALM’s ability to sense and induce membrane curvature^16,17^ has been associated with the insertion of AH0 into the membrane^16^ although it has never been directly demonstrated. The absence of AH0 in cells leads to the formation of larger flatter clathrin lattices that ultimately result in larger CCVs^6^, suggesting that AH0 is linked to the induction of membrane curvature.

In the characterization of model biomembranes, grazing incidence X-ray diffraction (GIXD) data provide in-plane molecular structure and lateral packing of phospholipid monolayers at fluid interfaces, while specular neutron reflectometry (NR) measurements can yield information about out-of-plane molecular structure and composition. It is important to note that, unlike X-rays, neutrons are non-destructive and highly penetrating, thus allowing measurements with sub-nanometer resolution at energies corresponding to thermal fluctuations, without inducing radiation damage. Moreover, as neutrons interact very differently with hydrogen (^1^H) and deuterium (^2^H), isotopic substitution can highlight structural differences in specific regions of interest, including the presence of water molecules as well as bound protein molecules. Solid-state nuclear magnetic resonance (ssNMR) spectroscopy can provide information about structure, dynamics, and molecular interactions with atomic-level resolution ^18,19^. This site-selective analytical technique was used to probe membrane-protein interactions, in a chemical environment mimicking *in-vivo* conditions, through ^1^H, ^13^C, and ^31^P isotropic chemical shift (δ_iso_), ^31^P chemical shift anisotropy (CSA) and longitudinal relaxation (T_1_) analyses. Unlike electron spin resonance (ESR) spectroscopy techniques, which require the addition of large planar hydrophobic probes at positions along the length of a supposed membrane inserting helix, ssNMR is carried out on native proteins. ssNMR is therefore not subject to the major caveats and issues from which ESR can suffer.

To shed further light on the role of CALM as a driver of membrane curvature, the interactions between the protein and *in vitro* biomimetic PtdIns(4,5)P_2_ presenting model plasma membranes of increasing complexity were characterized at room temperature and under physiological-mimetic conditions using a combination of GIXD, NR and ssNMR, along with other biophysical techniques. These studies aim at recreating important elements of the binding mechanism of CALM by simplifying the system using appropriate minimal model membranes, including lipid monolayers and solid-supported bilayers. In this regard, to assess membrane insertion of the CALM’s AH0, the wild-type ANTH domain of the protein (hereinafter CALM wild type, CALM_wt_) and a mutant missing the AH0 (CALM_ΔAH0_) were compared.

## Results and Discussion

### Phospholipid Langmuir monolayers as models of the inner leaflet of the plasma membrane

The inner leaflet of the mammalian plasma membrane is characterised by the presence of phosphatidylcholine (PC) and phosphatidylethanolamine (PE)^20–22^ as well as phosphatidylinositol 4,5-bisphosphate, PtdIns(4,5)P_2_ (∼1-2 mol %) ^23–26^. The existence of a non-specific, electrostatic interaction between CALM and the negatively charged inositol-biphosphate ring of PtdIns(4,5)P_2_ was initially investigated using an *in vitro* model system based on a Langmuir lipid monolayer at the air/buffer interface. For this purpose, a lipid mixture containing 1, 2-Dipalmitoyl-sn-Glycero-3-Phosphocoline (DPPC), 1, 2-Dipalmitoyl-sn-Glycero-3--Phosphoethanolamine (DPPE) and PtdIns(4,5)P_2_, in a molar ratio of 7:2:1, was chosen (molecular structures shown in ***Figure S1***), based on a similar composition of phospholipid headgroups previously used in the study of CALM^16^. DPPC and DPPE were selected for their stability at the air/water interface, as their saturated palmitoyl chains prevent oxidation^27,28^.

The phase behaviour of PtdIns(4,5)P_2_-enriched model monolayers was evaluated by the surface pressure (Π) - area per molecule (A) isotherm reported in ***Figure 1A***. The lateral compressibility of the monolayer, which indicates its mechanical resistance against dilational deformation, can be determined from the slope of the Π-A isotherm. This monolayer intrinsic property, commonly expressed in terms of the compressional elastic modulus (C_s_ ^-1^), is usually linked to the membrane fluidity and thus can be compared against the molecular area (***Figure 1C***). Two phospholipid packing densities (indicated by arrows in ***Figure 1A***) at surface pressures of 15 mN/m and 25 mN/m were chosen to explore the effect of lipid lateral packing on CALM binding. These values represent regions of coexisting non-miscible fluid/condensed phases with relatively low and high compressibility modulus, respectively, based on the increase in the linear dependence between C_s_^-1^ and Π shown in ***Figure 1B*** (from C_s_ ^-1^ ≈ 2Π to ≈ 6Π). The existence of DPPC:PtdIns(4,5)P_2_ immiscible phases in a lipid mixture monolayer was previously reported ^29^.

**Figure 1.**
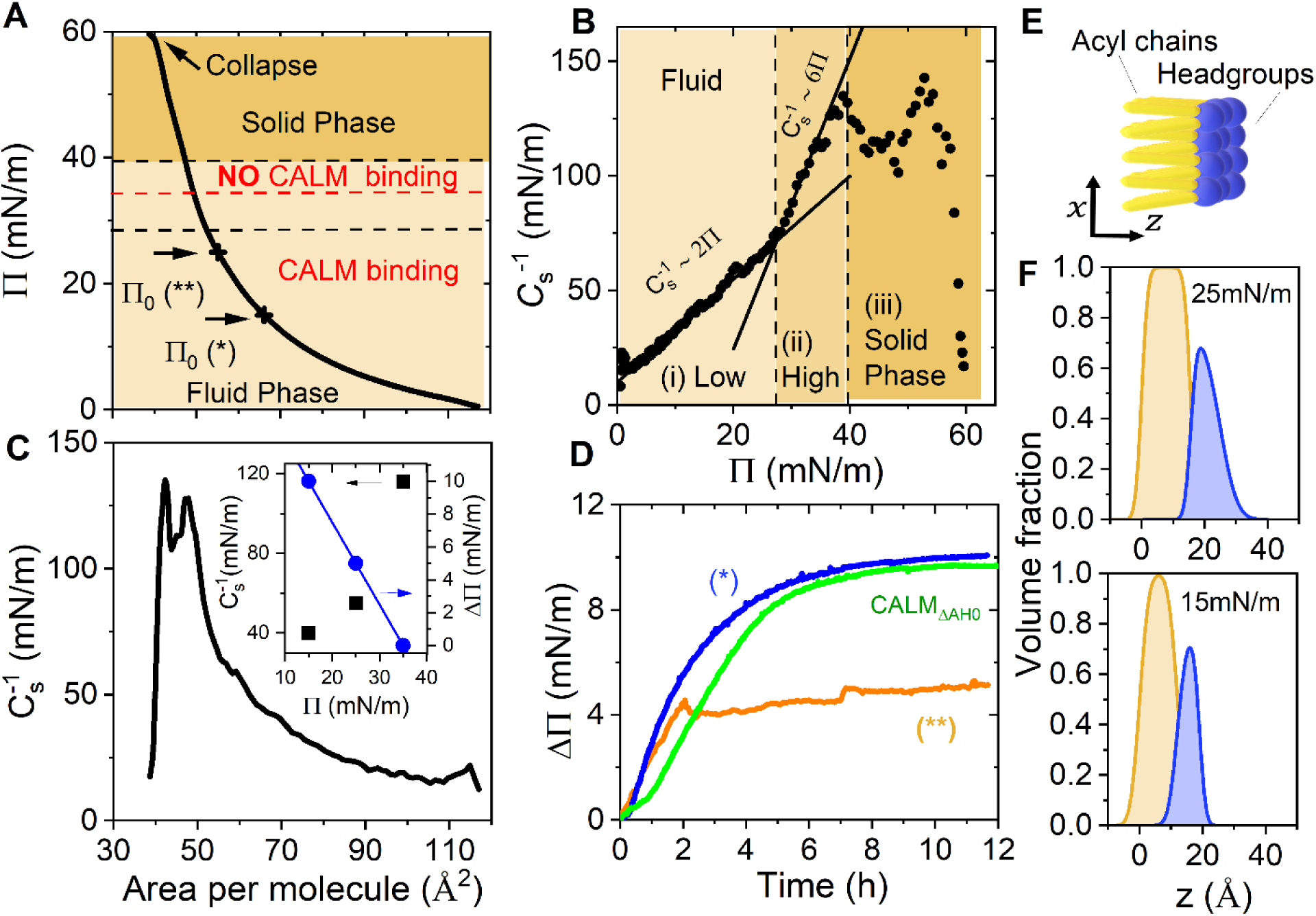
CALM interaction with inner leaflet of the plasma membrane. **(A)** Π − α isotherm of DPPC:DPPE:PtdIns(4,5)P_2_ lipid monolayer. The arrows indicate the values of Π selected for NR studies. Lateral compressibility moduli (*C* ^−1^) as a function of Π **(B)** and A **(C)**. **(D)** Increase in surface pressure ΔΠ upon the injection of CALM_wt_ or CALM_ΔAH0_ in the bulk phase at Π_0_ = 15 ^mN·m-1^ and 25 ^mN·m-1^. **(E)** Cartoon of a lipid monolayer representing also the stratified layer model (with headgroup and tail regions) used to fit the NR data. **(F)** Volume fraction profiles of the lipid monolayers derived from the fitting of the NR distribution profiles plotted in **Figure S2**. Inset in **(C)** shows how the increase in *C* ^−1^ (black squares) with Π can be correlated with ΔΠ (blue circles) decrease corresponding to CALM_wt_ binding.

Phospholipid structure and composition orthogonal to the plane of the leaflet at two different lateral pressures, denoted as Π_0_, were determined by NR. Analysis of NR data collected for hydrogenous lipid monolayers in different isotopic contrasts is shown in ***Figure S2***. PtdIns(4,5)P_2_-enriched monolayers could be effectively modelled as two layers (***Figure 1E*** and ***Materials and Methods***) corresponding to (i) the hydrophobic aliphatic lipid tails protruding into air, and (ii) the hydrophilic lipid headgroups in contact with the buffer. The lipid area per molecule obtained from NR was consistent with values obtained from the Π-A isotherm, as shown in ***Table S2***. From the neutron scattering length density (SLD) distribution (***Figure S2***), the volume fraction of the different monolayer components was determined and plotted in ***Figure 1F***. The thickness of the lipid headgroups layer was 8 ± 1 Å, and 9 ± 1 Å, with a solvent penetration of 35 ± 1 % and 24 ± 1%, respectively for 15 and 25 mN/m. Moreover, as expected, the increase in surface pressure led to an increase in the thickness of the layer representing the aliphatic chains (from 12 ± 1 Å to 14 ± 1 Å).

### Interaction of CALM with the plasma membrane: the role of lipid packing

At the surface pressure Π_0_ of 15 mN/m, the resulting lipid packing area per molecule of 65 Å^2^ is close to that expected for a fluid-like lipid bilayer^30^. Subsequent injection into the bulk buffer underneath the monolayer of CALM to a final concentration of 5 µM gave rise to an immediate and rapid increase in surface pressure, followed by a slower increase until a plateau was reached (ΔΠ = Π - Π_0_ = 9.5 ± 0.5 mN/m) (***Figure 1D***). This observation suggests the favourable recruitment of CALM into fluid membranes with high compressibility (inset in ***Figure 1B***). Consequently, CALM binding should exert a net force on the lipid surrounding, which reacts with an equivalent compression force yielding an increase in the surface pressure. In the case of a denser packed monolayer and lower fluidity (55 Å^2^, at Π_0_ = 25 mN/m), the increment in pressure was lower (ΔΠ = 5 ± 0.5 mN/m) reflecting a reduced recruitment of CALM_wt_. This can be rationalised with the tightly packed and highly ordered lipid monolayer. The resulting lower compressibility could prevent CALM from properly accessing the lipid surface, thus inhibiting binding. These results strongly support the fact that CALM_wt_ interaction is strongly influenced by the lipid packing of the membrane reflected by the compressibility modulus increase. Favourable binding of CALM is thus observed in the low compressibility region (***Figure 1B***).

NR was used in the following to disentangle the coverage of CALM_wt_ interacting with the membrane. No further CALM binding was observed at values of Π > 35 mN/m that can be rationalised by the increase in the compressibility modulus (inset in ***Figure 1C***). The reduction in ΔΠ with the increase in Π_0_, also reported in the inset of ***Figure 1C***, may be due to a progressive decrease in the number of proteins binding to the monolayer or a reduced extent of protein insertion into the monolayer, driven by steric constraints due the diminishing availability of free area. This hypothesis was explored through further NR experiments.

### CALM orientation and binding preference for PtdIns(4,5)P_2_ monolayer headgroups

To evaluate the role of lipid lateral density on CALM_wt_ binding, NR experiments were performed to determine the structures of CALM_wt_ bound to PtdIns(4,5)P_2_ enriched monolayers. ***Figures S2D*** and ***S2J*** show the reflectivity profiles in three isotopic contrasts after the binding of CALM_wt_ to monolayers at 15 and 25 mN/m. The obtained reflectivity curves before and after protein injection at 15 mN/m showed significant differences (***Figure S2D***), compared to the ones at 25 mN/m whose differences were almost negligible (***Figure S2J***). This aligns with surface pressure measurements showing the CALM preference to high compressible, fluid monolayers.

The volume fraction of CALM_wt_ in the orthogonal direction to the membrane plane (See scheme in ***Figure 2E***) derived from the NR data collected on PtdIns(4,5)P_2_ monolayer at 15 mN/m shows that CALM_wt_ is positioned beneath the membrane, exhibiting limited insertion only at the level of lipid headgroups (≈ 13 ± 1%) and a negligible presence in the acyl chain region (see ***Figure 2A*** and ***Table S3***). Conversely, at 25 mN/m CALM_wt_ does not display observable insertion into the membrane. ***Figure 2A*** shows that only 2% of the protein is situated beneath the monolayer. This observation implies that reduced surface accessibility, due to a denser lipid arrangement, hinders CALM_wt_ from integrating into the hydrophobic core of the membrane and reaching the lipid head groups on the surface. As many protein binding interactions occur at the lipid headgroup interface, a reduction in available surface area can hinder these binding events due to steric hindrance ^31^. Moreover, the orientation of the phosphorylated inositol ring of PtdIns(4,5)P_2_, dependent on the lipid packing fraction, might impact CALM preferential recruitment. As Π_0_ is reduced from 25 to 15mN/m, the inositol ring moves closer to the membrane, *i.e.*, it positions itself towards a parallel orientation with respect to the monolayer ^29^. This orientation is hypothesised to facilitate CALM binding.

**Figure 2.**
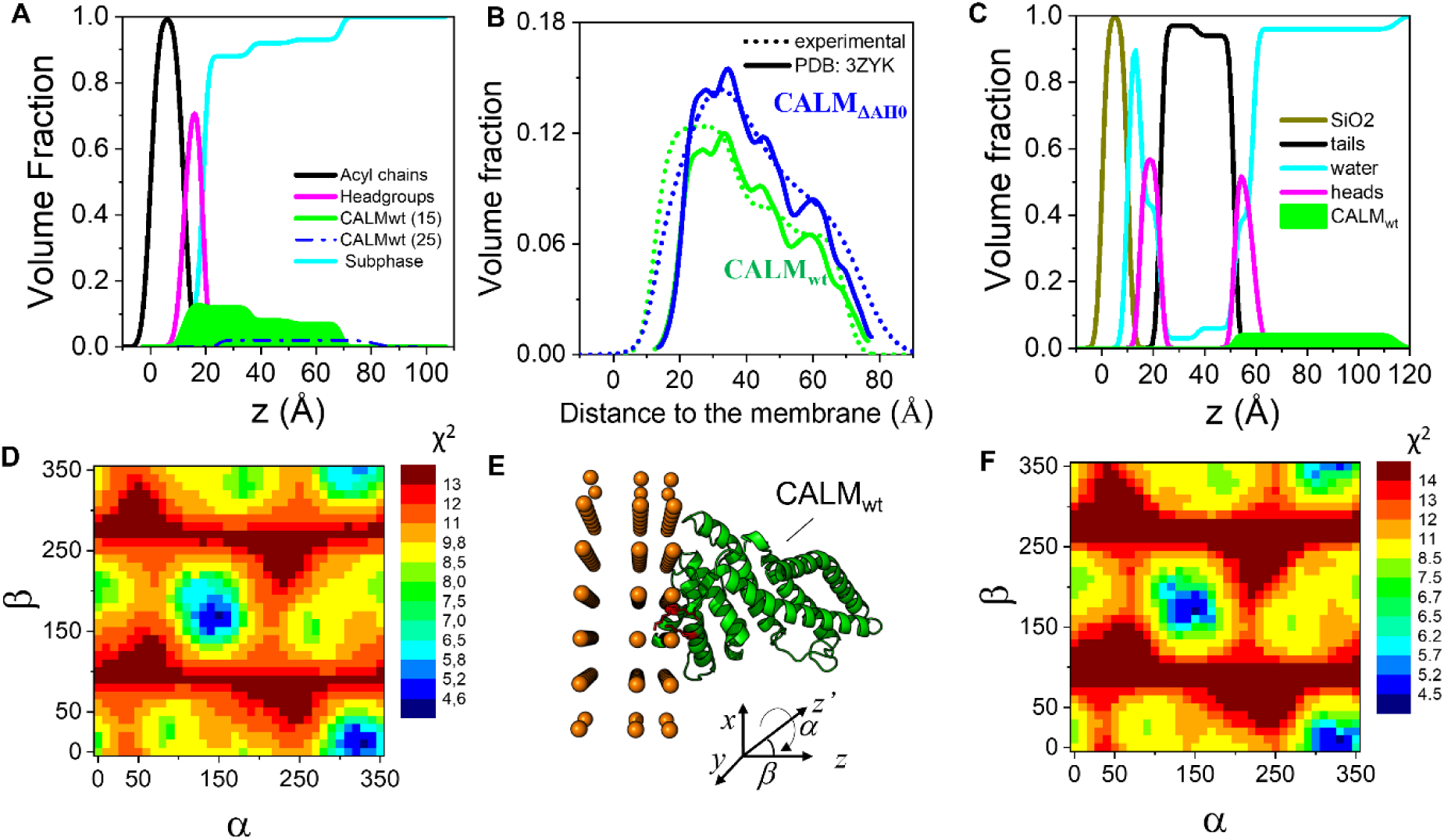
High resolution NR analysis to obtain the preferred orientation of CALM bound to PtdIns(4,5)P2 enriched mono- and bilayers. Experimental data and fits reported in SI (Figures S2 and 3) In-plane averaged spatial distributions profiles of CALM inserted in monolayers (A) and solid-supported bilayers (C) respectively, expressed as volume fractions (, of lipid components and CALM_wt/(AH0_ in the direction orthogonal to the plane of the interface. (B) Experimental volume fraction profiles compared to theoretical ones calculated for CALM ANTH domain (PDB: 3ZYK, residues 19-288) conformation based on the best possible orientations for CALM_wt_ and CALM_ΔAH0_ bound to lipid monolayers. Colormaps showing the orientation of CALM_wt_ (A) and CALM_ΔAH0_ (C) that can best fit the experimental data (see SI Figure S4). (E) Molecular cartoon representation of the best orientation (α=330°, β=10°) of CALM. The PtdIns(4,5)P_2_ binding site, which is depicted in red, points towards the membrane. The orange spheres indicate the interfaces between the slabs, i.e., air/tails-layer, tails-layer/headgroups-layer, headgroups-layer/bulk.

To further explore the association of CALM with membranes, solid-supported lipid bilayers (SLBs) with 1,2-dioleoyl-sn-glycero-3-phosphocholine (DOPC), 1,2-dioleoyl-sn-glycero-3-phosphoethanolamine (DOPE) and enriched in PtdIns(4,5)P_2_ (7:2:1 molar ratio) were studied. SLBs were prepared by liposome adsorption and fusion onto a solid substrate (see ***Materials and Methods*** section). The bilayer described by NR would incorporate a symmetrical structure with solvated head groups of an inner leaflet, in contact with the solid support, and of an outer leaflet, in contact with the bulk phase. The acyl tail layers had a similar thickness of 15 Å (***Figure S3*** and ***Table S4***). The characterised bilayer had minimum water content in the acyl tail region thus confirming almost full lipid coverage (higher than 89%). Bilayer integrity was conserved after both hydrogenous (h-CALM_wt,_ ***Figure 2C***) and fully deuterated protein (d-CALM_wt_, ***Figure S3***) binding. By assuming that the structure of the membrane and its bound CALM are both invariant under isotopic substitution, reflectivity data changes on addition of CALM are compatible with the partition of CALM into the membrane’s outer headgroups region (≈ 5 ± 1% volume occupancy for h-CALM_wt_, and 3 ± 1% for d-CALM_wt_). The best model derived from the NR analysis incorporates an extra layer that would account for protein present outside the bilayer, with a thickness of 56 ± 1Å and a volume fraction of 7.0 ± 0.2% and 4 ± 1% for h- and d-CALM_wt_, respectively, as shown by the component volume fraction profiles plotted in ***Figure 2C***. These results agree rather well with previous NR experiments performed on monolayers at 15 mN/m, in which CALM only bound to the headgroups and expanded to the bulk with a similar volume fraction close to the membrane (***Figure 2A***). The area per lipid molecule at 15 mN/m of the monolayer matches with the one of the SLBs as reported by NR analysis, making those systems comparable.

The initial NR data analysis of membrane-bound CALM_wt_ yielded a low-resolution volume fraction profile derived from a slab model of constant scattering length density that is inherently linked to the size of a protein tertiary structure (***Figure 2A***). To enhance the precision of NR data analysis, atomistic protein models were employed to infer CALM_wt_ preferred orientation on the membrane (***Figure S4*** and ***S5***). SLD profiles were calculated from atomic models of CALM (based on energy minimised crystal structures from PDB 3ZYK^6^) from amino acids 19 to 288 and lacking AH0, since its position upon binding remains unknown. An ensemble of SLD profiles from 1296 different orientations was generated with rigid-body rotations around the center of mass of the protein by 10° increments in the Euler angles *α* and *β* (following an x-y-z extrinsic rotation scheme; the third Euler angle *γ* is not relevant as the SLD is plane-averaged in the z-direction). Following each rotation, the position along the axis orthogonal to the plane of the membrane, was optimised by fitting the resultant theoretical and experimental SLD to obtain an optimised protein volume fraction (***Figure 2B***). The optimum orientation of the protein structure was identified at the complementary pair of Euler angles (α, β) = (330°, 10°) and (150°, 170°). The CALM_wt_ is found positioned in contact with the lipid head groups, in a biologically appropriate orientation. In particular, the residues inserted in the headgroups layer include Lysines 25, 38 and 40, that directly interact with PtdIns(4,5)P_2_ (***Figure 2E***). Importantly, the resultant total surface coverage of 12% is similar to the coverage obtained by Miller et al. with PtdIns(4,5)P_2_ containing liposomes (16%)^16^.

To gain further information at atomic-level resolution about the binding of CALM_wt_ to the plasma membrane model, lipid bilayers composed of 1,2-dioleoyl-sn-glycero-3-phosphocholine (DOPC) and PtdIns(4,5)P_2_ (***Figure S1***) were prepared and characterized by solid-state NMR (ssNMR) in the presence and absence of the CALM_wt_ protein.

The ^1^H, ^13^C and ^31^P direct-excitation spectra of DOPC (with and without CALM_wt_ protein) depicted in ***Figure 3*** are largely dominated by the signals of DOPC lipid molecules, which are the most abundant (DOPC:PtdIns(4,5)P_2_ molar ratio of 97:3, determined from ^31^P integral). The ^31^P NMR spectrum in the absence of CALM_wt_ (***Figure 3A***) clearly shows the phosphate signal of DOPC at 0.9 ppm, alongside with three other less intense resonances at 0.3, 1.0 and 2.1 ppm assigned to the diester phosphate, and to the P5 and P4 inositol phosphate groups of PtdIns(4,5)P_2_, respectively. These chemical shift values are in good agreement with previous reports for membrane bilayers ^32,33^. After protein incubation, a clear shift of the PtdIns(4,5)P_2_ phosphate (*δ*_iso_) was observed from 2.1 (P4) and 1.0 (P5) ppm to 2.7 and 1.6 ppm, respectively (***Figure 3A***). The shift towards lower frequencies suggests a more shielded environment, which is consistent with a decrease in the negative surface charge of the bilayer induced by the positively charged protein residues (K28, K38 and K40^13^) upon binding, as previously observed in similar systems ^34^. The ^31^P observed chemical shift perturbation (≈ 0.6 ppm) is relatively small compared to other biomolecular systems, however, it fits within the expected values. Typically, ^31^P NMR chemical shift deviations in phospholipids interacting with proteins fall within the range of 0.1 to 5 ppm, depending on the interaction type and experimental conditions ^35–37^. These shifts reflect changes in the phosphate group’s electronic environment, influenced by electrostatic interactions with protein residues, bilayer structural alterations, or lipid packing variations. However, hydrogen bonding or ionic interactions, such as those involving CALM and PtdIns(4,5)P_2_, often result in shifts near the lower end of this range (0.1–0.5 ppm) ^35,38,39^.

**Figure 3.**
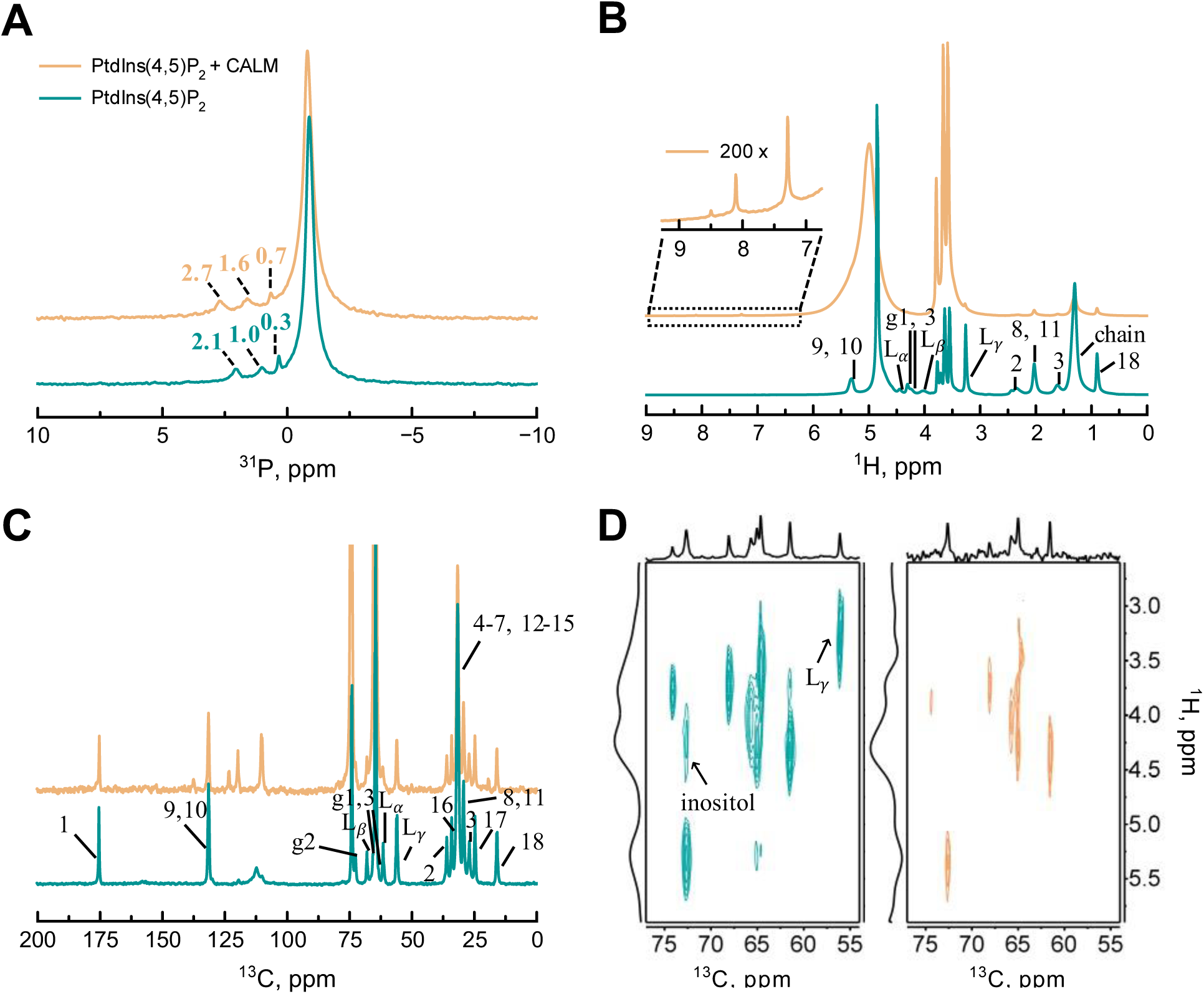
Solid-state NMR spectra: 1D direct excitation of ^31^P (A), ^1^H (B),^13^C (C) and 2D ^13^C-{^1^H} CPMAS HETCOR (D). The inset of ^1^H spectra (B) show the presence of peaks at 7.3, 8.1 and 8.5 ppm, characteristic of protein amide protons. The cross-peaks of the PtdIns(4,5)P_2_ inositol and DOPC choline terminals (denoted by an arrow), in the 2D ^13^C-{^1^H} CPMAS HETCOR spectrum (D), disappeared after the binding of CALM_wt_. All the spectra were acquired at 290 K using a MAS speed of 8 kHz.

The insertion of PtdIns(4,5)P_2_ in the bilayer leads to broadening of the DOPC signal (***Figure S6***). The broadening of the DOPC signals is higher after the binding of CALM_wt_ (***Figure S7*** and ***Table S6***). It is noted that when CALM_wt_ was incubated with DOPC liposomes lacking PtdIns(4,5)P_2_ no changes were observed and the signal width is comparable to DOPC without protein, demonstrating that the docking of CALM_wt_ to the membrane is happening only when PtdIns(4,5)P_2_ lipids are present. Spectral deconvolution of DOPC/PtdIns(4,5)P_2_ liposomes in the presence and absence of the CALM_wt_ is shown in ***Figure S7*.**

One common way in which peripheral proteins and peptides reach their biologically active membrane-bound state, involves initial electrostatic binding via positively charged domains to negatively charged lipid bilayers followed by membrane insertion of hydrophobic domains^34^. This type of binding action is consistent with the results observed for the binding of CALM_wt_, where ^31^P *δ*_iso_ shows a decrease in the negative charge of the bilayer induced by the interaction with the positively charged residues in the binding pocket of the protein ^13,16^. Moreover, the results from NR corroborate this hypothesis by demonstrating that the AH0 helix of CALM_wt_ is inserted into the bilayer upon binding. The ^31^P signal of DOPC exhibits small *δ*_iso_ variation (Δ ^31^P ≈ 0.1 ppm) and broadening of the signals in the presence of the protein. This broadening is typically associated with a lower mobility, which is associated with increased membrane rigidity.

***Figure 3B* and *3C*** depict the ^1^H and ^13^C NMR spectra, which are largely dominated by the signals of DOPC. Therefore, to avoid misleading assignments, only the signals of DOPC lipid molecules were assigned in the ^1^H and ^13^C direct-excitation NMR spectrum (see ***Figure S8*** for chemical group legend). The assigned signals are in good agreement with previous works ^40–47^. Small signals, which are present as shoulders in the dominant DOPC peaks, are attributed to the PtdIns(4,5)P_2_ lipids. The ^1^H NMR spectrum (Figure 3B) shows the presence of peaks at 7.3, 8.1 and 8.5 ppm, after protein incubation, which are characteristic of amide protons^48^.

2D ^13^C-{^1^H} CPHETCOR spectra was obtained in the presence and absence of the protein (Figure 3D). After the binding of CALM_wt_ to the membrane, major changes in the cross-peaks correlation area around 55 – 75 ppm (^13^C dimension) were observed (Figure 3D). This was expected considering that the phospholipid headgroup signals appear mostly in this area, *i.e.*, ^13^C signals of the phospholipid glycerol backbone and polar groups. Notably, the PtdIns(4,5)P_2_ inositol and the choline terminal CH_3_’s (L*γ*) cross-peak signals (marked by an arrow in Figure 3D) completely disappear after protein incubation. The remaining lipid cross-peak signals also exhibit some *δ*_iso_ changes with the most prevalent shifts occurring at the headgroup level as can be observed by the *δ*_iso_ variation plot in Figure 4. These results indicate that the interaction of CALM_wt_ and the lipids occurs at the headgroup level, confirming the results observed from NR. *δ*_iso_ changes are also observed in the aliphatic chains of the lipids. This *δ*_iso_ variation coupled to the broadening of the ^31^P DOPC signals, after protein incubation, indicate a change in the plasmatic membrane packing which is induced by the anchoring of the AH0 helix in the bilayer, as demonstrated by NR.

**Figure 4.**
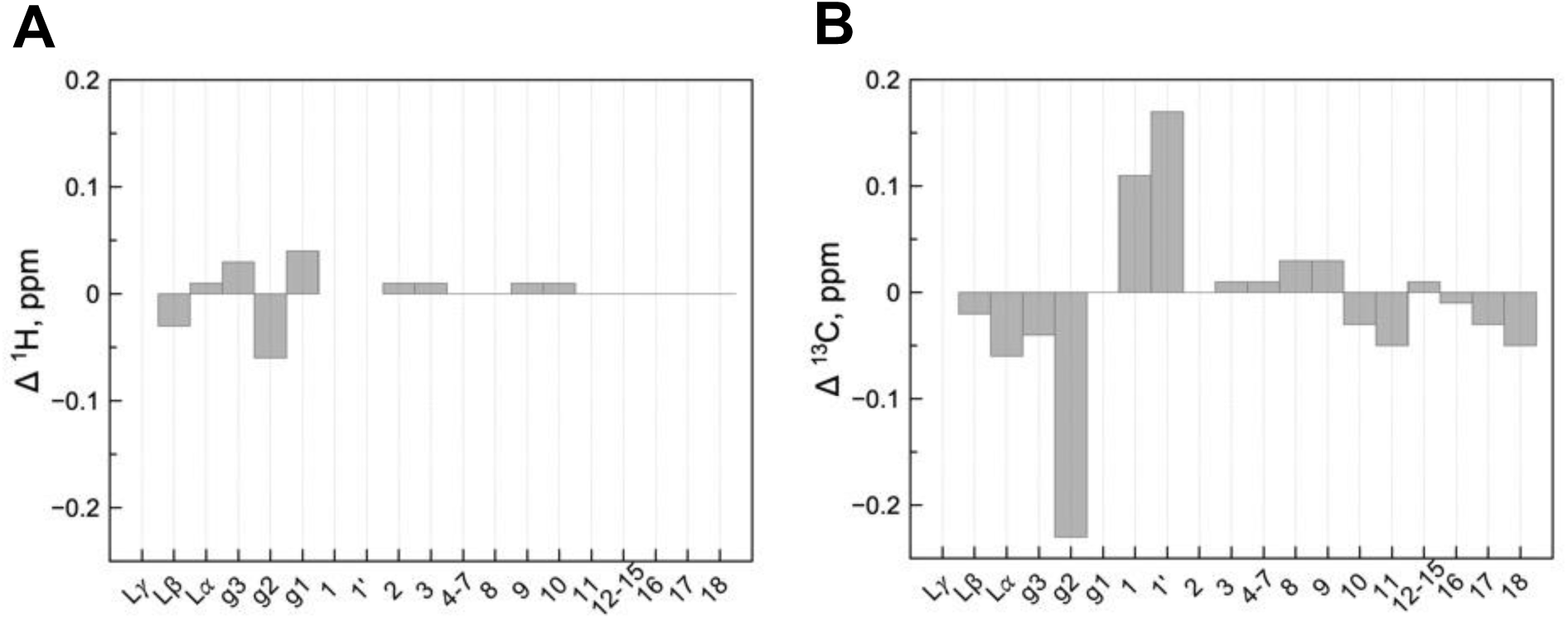
Chemical shift variation plot of ^1^H (A) and ^13^C (B). The data of this plot was derived from the CS difference of ^13^C-{^1^H} CPHETCOR experiments before and after CALM_wt_ incubation.

CSA and T_1_ relaxation measurements were performed to assess the overall dynamics of the system in the presence and absence of the protein. Longitudinal relaxation T_1_ times for the phosphate ^31^P lipid signals are reported in ***Table S7***. ^31^P T_1_’s in the protein-free liposomes exhibit values in the range of those reported in the literature at room temperature for phospholipids ^36,49,50^. The encountered short values are the reflection of a dynamic environment leading to strong modulation of CSA interaction, becoming a very efficient relaxation mechanism. ^31^P CSA (see ***Supporting Information*** for additional CSA details) has been proposed as main interaction-driving relaxation in phospholipids at high magnetic fields ^51^. Therefore, phosphate groups in DOPC and PtdIns(4,5)P_2_ with larger *δ*_CSA_ values (***Table S8***) have shorter relaxation times (***Table S7***). After protein incubation, ^31^P T_1_ relaxation times for PtdIns(4,5)P_2_ inositol phosphates become shorter (***Table S7***). This reduction is associated with a reduction in mobility throughout the membrane upon CALM_wt_ binding as chemical shift analysis and NR data suggest. The slower dynamics of the PtdIns(4,5)P_2_ head groups induces a less effective averaging of the CSA ^50,52–55^. The reduction in ^31^P longitudinal relaxation times in for PtdIns(4,5)P_2_ is the result of the direct interaction of this phospholipids with CALM_wt_ as represented in Figure 7.

Additionally, CSA tensor parameters were determined by acquiring ^31^P direct-excitation spectra at variable spinning speeds (***Figure S9***). The ^31^P CSA values reveal an increase in *δ*_CSA_ parameter of all the phosphates of PtdIns(4,5)P_2_ lipids after protein incubation (***Table S8***). PtdIns(4,5)P_2_ increased CSA is associated with a decrease in the mobility of the lipid upon binding of CALM_wt_.

### Assessing amphipathic helix 0 (AH0) insertion into lipid membrane

Amphipathic helices (AH), such as the CALM AH0 helix are known to interact with biological membranes, playing a role in various cellular processes^56–58^. Although AH0 is predicted to insert into the membrane^16^, the insertion depth and the nature of the interaction with the membrane had not been previously addressed. Langmuir tensiometry and NR were therefore used to analyse the interaction between the CALM mutant lacking the AH0 domain (CALM_ΔAH0_) and PtdIns(4,5)P_2_-enriched model monolayers.

The membrane binding affinity was assessed by measuring the surface pressure upon protein injection at Π_0_=15 mN/m. Both CALM_wt_ and CALM_ΔAH0_ rendered identical ΔΠ values following subphase injection (Figure 1D). This suggests that CALM recruitment at the membrane is not directly influenced by the AH0, but mainly relies on the electrostatic interaction between PtdIns(4,5)P_2_ and Lysines 28, 38 and 40^13^. Considering that the AH0 represents only 4% of the entire protein volume, the same slab model was used to fit CALM_wt_ and CALM_ΔAH0_ reflectivity data (see scheme in Figure 2E). The resultant analysis revealed comparable total extensions for CALM_wt_ and CALM_ΔAH0_ (57±4 Å and 63±4 Å, respectively) as well as similar volume fraction percentages underneath the monolayer (12% for CALM_wt_ and 15% for CALM_ΔAH0_) as shown in Figure 2B.

Further calculations of NR profiles were performed using rotated structures of CALM (based on PDB 3ZYK) and, the different calculated orientations of CALM_ΔAH0_ with respect to the monolayer were compared to the experimental NR data. This new refinement returned the same pair of Euler angles found for the CALM_wt_ structure, thus suggesting that the orientation is independent from the presence of AH0 (see Figure 2B). Though, the volume fraction percentage of CALM_ΔAH0_ inserted in the headgroups-layer was found to be only 5 ± 1%, with respect to the 13 ± 1% of CALM_wt_, coherent with the presence of fewer amino acid residues present in the membrane due to the absence of AH0 (***Table S3***).

Based on the identical structures derived from NR analysis of CALM_wt_ and CALM_ΔAH0_ binding, we propose that the AH0 domain of CALM fully inserts within the headgroup layer. This allows the calculation of CALM:PtdIns(4,5)P_2_ stoichiometry, as well as the protein volume inserted in the membrane per lipid molecule. Considering that PtdIns(4,5)P_2_ is 10% in molecules of the total lipid composition, we estimate that each CALM_wt_ molecule interacts with a cluster of 4-5 PtdIns(4,5)P_2_ molecules. This binding of the CALM’s ANTH domain to PtdIns(4,5)P_2_ clusters aligns with findings from coarse-grained molecular dynamics simulations employing variations of the terminal amphipathic helix^59^.

The roughness parameter, defined as the boundary between the peripheral CALM region and the lipid headgroup region, notably increases in the absence of the AH0 domain (8 Å compared to 3 Å in CALM_wt_, ***Table S2***), which would imply a potentially diminished protein-membrane interaction strength. This could be attributed to AH0’s putative function as an anchor for protein-membrane attachment.

### Effect of CALM on the in-plane structure of PtdIns(4,5)P_2_ enriched lipid monolayer would promote membrane bending in bilayers

To investigate the impact of CALM interaction on the packing of lipid aliphatic chains, grazing incidence X-ray diffraction (GIXD) profiles of PtdIns(4,5)P_2_-enriched monolayers were recorded using synchrotron radiation. GIXD is particularly effective for providing molecular-scale insights into the in-plane packing of aliphatic chains, as X-rays at a grazing angle penetrate deeply, on the order of tens of Ångstroms, into the air/water interface ^60^.

The observed Bragg peaks in the GIXD data were interpreted as being primarily due to the membrane acyl chains. ***Figure 5A* and *5C*** show the GIXD contour plots of the scattered intensity as a function of q_xy_ and q_z_ for a DPPC:DPPE:PtdIns(4,5)P_2_ monolayer prior and after CALM_wt_ inserted at a surface pressure value of 15 mN/m, respectively, while Figure 5B shows the corresponding distribution of Bragg peaks. Bragg rod analyses of the PtdIns(4,5)P_2_ enriched monolayer in the absence and presence of CALM_wt_ are shown in ***Figure S10***. As expected, the experiments did not show diffraction from the head groups of phospholipid molecules. Moreover, CALM_wt_ membrane binding was almost negligible at 25 mN/m and no difference in GIXD was observed before and after CALM_wt_ addition.

**Figure 5.**
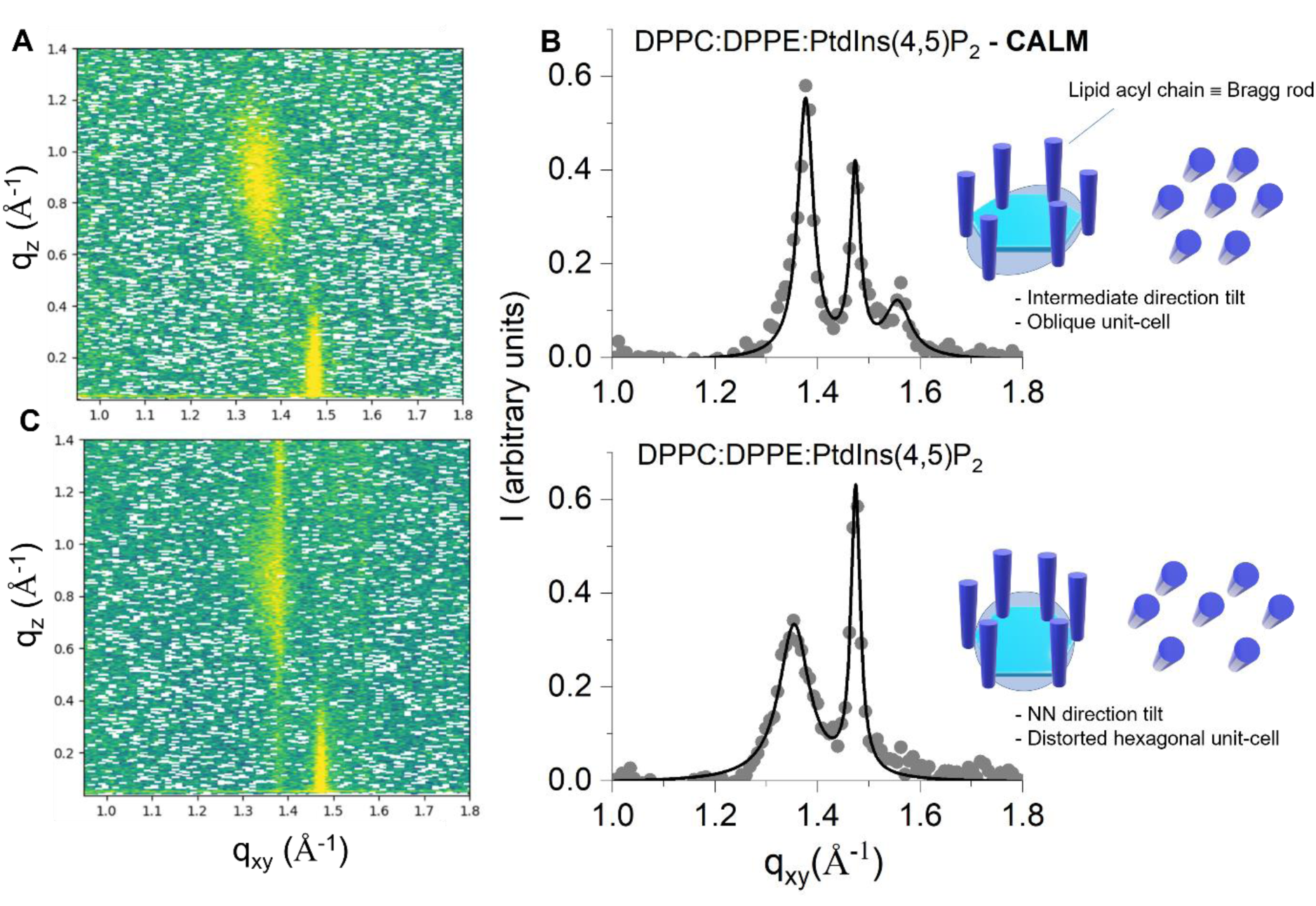
Inducing in-plane structural changes of lipid acyl chains by CALM binding to PtdIns(4,5)P2 enriched monolayers. GIXD intensity contours map for PtdIns(4,5)P2 enriched monolayers prior (A) and after CALM binding (C) at Π = 15 mN/m. (B) Distribution of Bragg peaks with q_xy_ obtained integrating over q_z_. Schematic side/top view representation of the lattice formed by DPPC/PE domains in the monolayer at the air/water interface. The position of the acyl chains is indicated.

In absence of CALM_wt_, two Bragg peaks (Figure 5B), centered at q_xy_= 1.35 Å^-1^ (at q_z_ = 0.85 Å^-1^) and q_xy_= 1.47 Å^-1^ (q_z_= 0.05 Å^-1^) were observed, which can be interpreted by means of a distorted 2D hexagonal unit cell with dimensions of |a| = |b| = 5.08 ± 0.05 Å , and γ = 114°, in which the acyl chains of the phospholipids are tilted towards their nearest neighbors (NN). The observed crystalline lattice is likely attributable to the presence of DPPC/PE-enriched condensed domains ^29^, consistent with the hexagonally distorted lattice (NN type) expected for a pure DPPC monolayer ^61^.

In the presence of CALM, three Bragg peaks, centered at q_xy_= 1.37 and 1.55 Å^-1^ (at q_z_ = 0.85 Å^-1^) and q_xy_= 1.47 Å^-1^ (q_z_ = 0.05 Å^-1^), were found and are consistent with an in-plane oblique lattice formed by the lipid chains. Bragg peaks were fitted using Lorentzian functions with the parameter reported in ***Table S5***. The lattice parameters corresponding to this oblique lattice were 4.99 ± 0.05 Å (a) and 4.66 ± 0.05 Å (b) with the angle (γ) between them equal to 114°. The intermediate tilt between NN and NNN (nearest and next-nearest neighbours, respectively) was 33.5 °, which was different to the value of 40 ° found in the absence of CALM. From these parameters a chain cross-sectional area A_0_ of 21 ± 1 Å^2^ was found, yielding an area per lipid molecule of A = 2 A_0_ / cos(φ) = 50 ± 1 Å^2^, with φ = 33.5° ± 0.5°, being the tilting angle calculated from the Bragg rod analysis (***Table S5***). There is a clear decrease in A (63 ± 2 Å^2^, in absence of CALM_wt_) that can be rationalized as an increasing of the molecular packing of the aliphatic chains produced by the monolayer compression due to the fraction of bound CALM_wt_. Despite only 12% of CALM_wt_ partially interacts with the monolayer headgroups, according to NR analysis (Figure 2A), these GIXD data suggest that CALM_wt_ binding induces a lateral redistribution of lipid packing throughout the membrane, extending beyond the immediate binding location. This has been observed also in other proteins bound to lipid monolayers^62^ and in molecular dynamics simulations of lipid - membrane proteins^63^. The hypothetical lateral propagation of area compression is probably facilitated through steric interactions among neighboring lipids, resulting in collective alterations in structure and long-distance rearrangement of lipid packing.

The modulation of the local lipid environment upon CALM binding, through changes in lipid packing observed in monolayers, suggests a mechanism by which CALM could promote membrane curvature in bilayers. These local changes in lipid packing could lead to a redistribution of mechanical stress within the plane of the membrane, potentially facilitating its deformation. In particular, differences in lipid packing between the bilayer’s leaflets—induced by CALM binding to one leaflet as reported in Figure 2C—could create an asymmetric environment favourable to curvature.

### Impact of CALM Binding on Lipid Bilayer Viscoelasticity, Local Stiffness, and Potential for Curvature Generation

To better understand the interaction between CALM and PtdIns(4,5)P_2_-enriched membranes, we employed Quartz Crystal Microbalance with Dissipation (QCM-D) monitoring. QCM-D is a highly sensitive analytical technique used to detect real-time changes in both mass and viscoelastic properties of solid-supported lipid bilayers. Upon adding CALM to previously assembled DOPC/DOPE/ PtdIns(4,5)P_2_ SLBs, a decrease in frequency (Δf) and an increase in dissipation (ΔD) were observed (Figure 6A-B). This frequency drop indicates the adsorption of additional material onto the surface of the quartz sensor, confirming that CALM adsorbs onto the bilayer surface. The simultaneous increase in dissipation suggests that CALM not only binds to the bilayer but also alters its mechanical properties. In fact, dissipation represents the ratio between the energy lost and that stored per oscillation cycle and evaluates all sources of energy dissipation in the system. When a viscoelastic film adheres to the quartz crystal, it deforms with each oscillation, leading to high dissipation values. In contrast, rigid material closely follows the crystal’s oscillation without any apparent deformation, resulting in low dissipation, typically below 1 ppm. As CALM binds, the dissipation increases slightly over time until it reaches a steady-state value around 2 ppm. This is indicative of a frictional component within the mechanical response of the deposited SLB, which reflects chains in the SLB structure. It is clear that CALM binding enhances the viscoelasticity of the lipid bilayer, rendering it more deformable. This effect likely arises from an increase in structural flexibility induced by CALM, which promotes rearrangement of the lipid molecules. These results align with GIXD data obtained from monolayers. These results collectively demonstrate that CALM’s adsorption onto the lipid bilayer not only increases the bilayer’s mass but also modulates its viscoelastic properties, implying that CALM induces a degree of structural rearrangement in the bilayer as it interacts with the PtdIns(4,5)P_2_-enriched surface.

**Figure 6.**
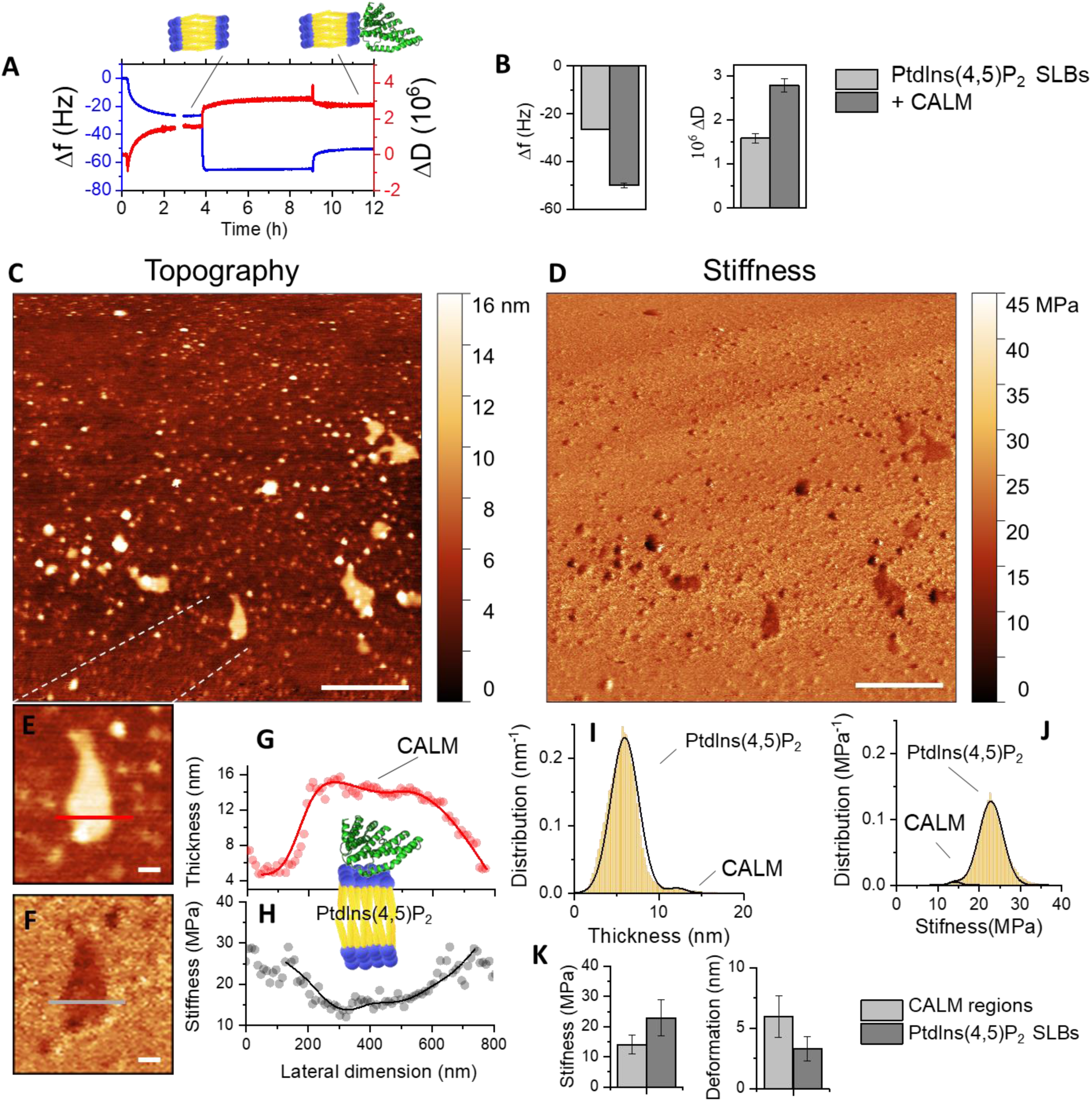
(A) Representative QCM-D measurements showing frequency (black) and energy dissipation (red) changes at the seventh overtone upon lipid bilayer formation and CALM addition.

In the following, we evaluate the local rigidity disruption induced by CALM binding within the membrane plane by using Fluid Phase Peak Force QNM mode AFM on DOPC/DOPE/ PtdIns(4,5)P_2_ SLBs. Prior to CALM binding, AFM topographic maps confirmed high lipid surface coverage, with minimal defects (< 2 % surface) and miscibility of the lipid components (***Figure S11***). Bright spots identified as PtdIns(4,5)P_2_ clusters, align with previous studies of biomimetic membranes^64,65^. Following CALM addition, an increase in the size and thickness of the bright spots is observed (Figure 6C), likely representing CALM binding to PtdIns(4,5)P_2_ clusters, as our previous structural NR studies indicated binding occurs at a ratio of 4-5 different PtdIns(4,5)P_2_ molecules per CALM molecule.

Peak Force QNM at very low loading forces (200 pN) allows simultaneous imaging and quantitative mechanical mapping with sufficient sensitivity to analyze solid-supported bilayers in the fluid phase, such as DOPC. This approach enabled us to acquire high-resolution maps of topography, stiffness, deformation, and adhesion force for DOPC lipid bilayers, highlighting regions of CALM binding (Figure 6D). Young’s modulus (E), which quantifies stiffness, was calculated using the Derjaguin-Muller-Toporov (DMT) model. Remarkably, the mechanical properties of PtdIns(4,5)P_2_-enriched bilayers were significantly altered by CALM binding, as indicated by AFM stiffness measurements (Figure 6D). In regions with CALM bound, as confirmed by the observed thickness increase (***Figures 6E*, *6G***) in agreement with NR data, stiffness decreased to 14.1 MPa, compared to 22 MPa in non-CALM regions (***Figures 6F, 6H***). This behavior was consistent across the SLBs, where a bimodal distribution of both thickness (Figure 6I) and stiffness (Figure 6J) emerged, corresponding to regions with and without CALM bound. This behavior was reproducible across other regions and AFM samples, yielding mean values presented in Figure 6K.

Remarkably, this reduction in stiffness confirms the scenario raised from QCM-D results and suggests that the presence of CALM induces local softening of the membrane, through changes in lipid packing and/or an increase in local compressibility. Such a decrease in stiffness may facilitate the membrane’s ability to deform and bend, thereby supporting the hypothesis that CALM acts as a curvature generator. A softer membrane will be able to accommodate curvature more readily, which is crucial for processes such as vesicle formation, where dynamic membrane invagination is essential. The mechanical environment in CALM-bound regions may thus predispose the membrane to curvatures that are not as readily achievable in stiffer, non-CALM regions. Furthermore, while the reduced stiffness indicates a more compliant membrane, it is important to recognize that this does not directly correlate with bending rigidity. The relationship between stiffness and curvature is complex; although lower stiffness may imply greater compliance, it does not inherently translate to increased bending without additional supporting data. Therefore, the observed mechanical alterations due to CALM binding provide valuable insights into how protein interactions can modulate lipid bilayer properties and facilitate curvature generation, further elucidating the role of CALM in membrane dynamics.

## Discussion and Conclusions

Clathrin-mediated endocytosis (CME) is the main mechanism by which eukaryotic cells selectively internalize proteins into the endosomal system. CALM, one of the most abundant clathrin adaptors, preferentially binds to PtdIns(4,5)P_2_ clusters in membranes. The insertion of its N-terminal amphipathic helix (AH0) into membranes has been proposed to either sense or induce membrane curvature. In this work, by using different biophysical methods on bottom-up, minimal, *in vitro* model biomembranes enriched in PtdIns(4,5)P_2_, CALM binding to model plasma membrane was investigated. The combination of interfacial tension measurements and NR results showed that the interaction between CALM_wt_ and an *in vitro* minimal model of the inner leaflet was dependent on lipid packing. An area per molecule typical of a fluid phase with high compressibility (low C_s_ ^-1^) promoted the favorable binding of CALM. This observation aligns with the binding affinity of CALM to SLBs that have similar molecular areas. However, CALM binding was hindered in a fluid phase monolayer with lower compressibility (larger values of C_s_ ^-1^ but still smaller than those characterizing a condensed phase), which is associated with a highly ordered lipid organization. High-resolution NR led to the determination of the protein position and orientation with respect to the membrane. The results obtained are consistent with *in vivo* data ^13,16^, as the PtdIns(4,5)P_2_ binding site on CALM (Lysines 28, 38 and 40) is oriented towards the membrane and partially inserted among the polar headgroups layer, thus allowing interaction with the PtdIns(4,5)P_2_ phosphate groups (Figure 7). Moreover, the NR analysis of lipid monolayers demonstrated that each CALM_wt_ molecule would bind to a cluster of 4 to 5 PtdIns(4,5)P_2_. The binding of a mutant lacking the AH0 helix (CALM_ΔAH0_), was also investigated. The NR analysis of both CALM_wt_ and CALM_ΔAH0_ bound to lipid monolayers directly demonstrated that AH0 positions itself within the headgroups layer, therefore forming a possible wedge, which would induce membrane curvature, as well as stabilise protein docking to the membrane. In this regard, the GIXD and NR observed reduction of area per lipid molecule due to CALM binding could be the molecular mechanism to initiate membrane bending prior to clathrin binding.

**Figure 7.**
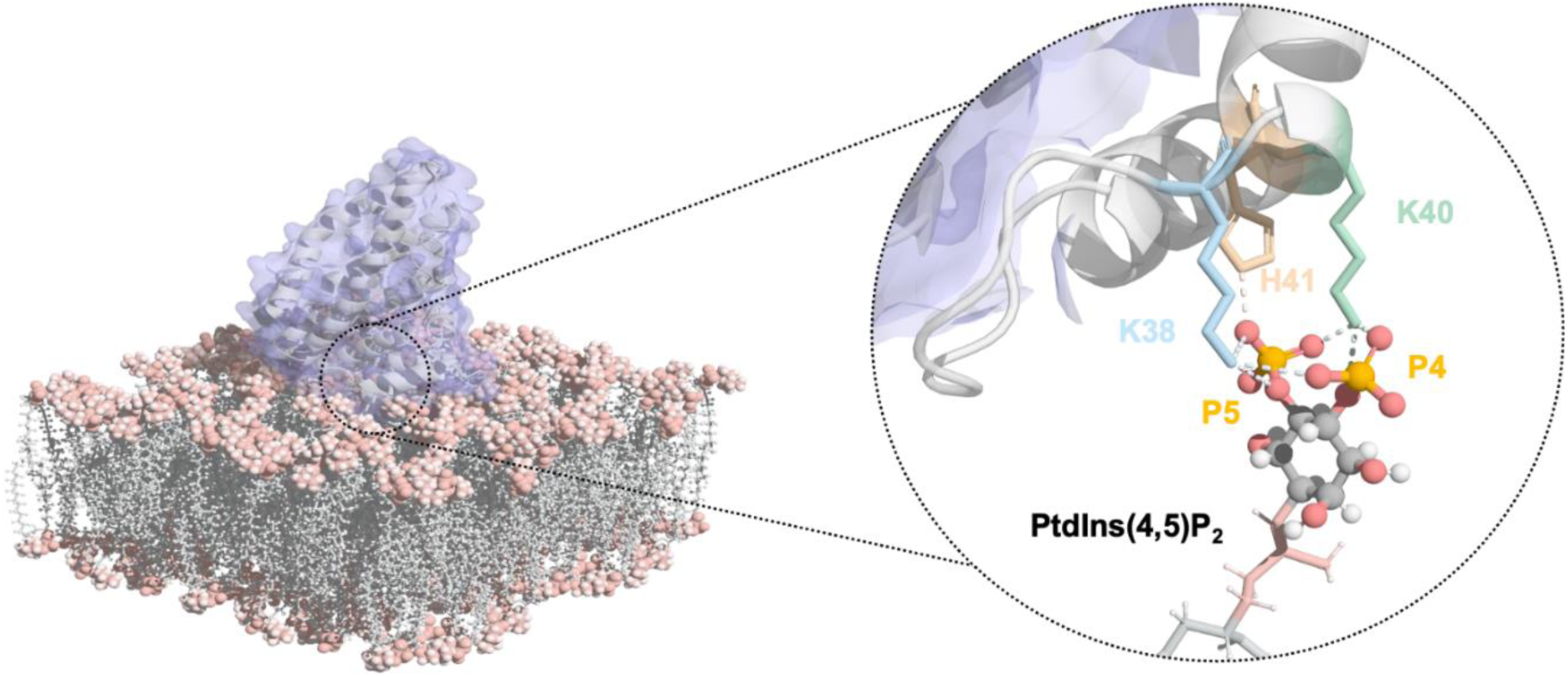
CALM folded domain is accommodated partly within the lipid membrane, directly interacting with PtdIns(4,5)P_2_ phosphates suggesting that LM’s amphiphilic N-terminal helix buries into the membrane. Structure of CALM interacting with a lipid bilayer enriched in PtdIns(4,5)P_2_ derived from the ssNMR and NR results showing in detail the interaction of CALM with the phosphates P5 and P4 from the inositol phosphate headgroup.

Our GIXD data show that CALM binding to PtdIns(4,5)P_2_-enriched monolayers induces tighter lipid packing, evidenced by a reduced area per lipid molecule. This structural compression likely creates a locally ordered and compressed leaflet, consistent with the hypothesis by Hossein and Deserno^66^, which suggests that differential stress, rather than composition asymmetry alone, dominates the lipid bilayer’s mechanical response to bending.

The increased viscoelasticity of the membrane observed through QCM-D further supports this interpretation. The accompanying rise in dissipation upon CALM adsorption indicates a significant frictional component and increased deformability within the bilayer. These findings suggest that CALM binding not only enhances lipid packing but also introduces greater flexibility and energy dissipation mechanisms into the membrane. This dual effect reflects an overall modulation of membrane mechanical properties that may facilitate structural rearrangements critical for dynamic processes such as curvature generation. The findings by Rickeard et al. complement this view, showing that asymmetric membranes exhibit altered stiffness governed by the more rigid leaflet, while interleaflet coupling introduces new dissipation contributions^67^. Such coupling-driven changes in membrane dynamics influence protein conformational changes, intermembrane interactions, and bending fluctuations. In our system, CALM-induced lipid rearrangements likely affect interleaflet coupling, as reflected in the observed increase in viscoelasticity and local membrane softening. This modulation may reduce the energetic barrier to curvature formation, promoting membrane deformability during processes such as endocytosis and trafficking.

The reduced stiffness observed through AFM Peak Force QNM corroborates this hypothesis, demonstrating that CALM binding induces local softening while enhancing lipid mobility. Collectively, the combination of GIXD, QCM-D, and AFM results reveals how CALM binding reorganizes the lipid bilayer, increasing its viscoelasticity and deformability while redistributing mechanical stress. These changes align with the models of Deserno and Rickeard et al., highlighting the critical role of lipid-driven mechanical properties in membrane remodelling and curvature generation ^66–68^.

Finally, ^31^P ssNMR spectroscopy revealed that upon binding to CALM the PtdIns(4,5)P_2_ P4 and P5 inositol signals shift to higher CS (*i.e.* shielded environments) which is typical of interactions with positively charged residues. This observation aligns with the anticipated binding domain of CALM (Figure 7), as proposed in earlier crystallographic studies, thereby providing additional evidence for its validity within an environment replicating *in vivo* conditions. Additionally, the DOPC phosphate ^31^P signal displayed significant broadening upon CALM_wt_ binding, suggesting an increase in membrane rigidity. The ^31^P CSA analyses revelaed an increase in *δ*_CSA_ for the P4 and P5 inositol signals of PtdIns(4,5)P_2_, accompanied by a significant *δ*_CSA_ increase in the broad component of DOPC. These findings suggest that binding effects are not localized but instead drive overall membrane remodelling through tighter lipid packing. These results are consistent with observations from Π-A isotherm, NR and GIXD, demonstrating increased lipid packing upon CALM binding. This phenomenon is attributed to the insertion of the AH0 domain into the membrane continuum. ^31^P CSA and T_1_ relaxation analyses further indicate an increase in rigidity of the membrane after biding of CALM_wt_.

In summary, our findings suggest a mechanism where CALM’s interaction with PtdIns(4,5)P_2_, leads to asymmetric lipid packing change relative to the membrane plane promoted by the insertion of amphipathic helix (AH0), thereby causing membrane bending.

## Materials and Methods

### Phospholipids and reagents

1,2-dipalmitoyl-sn-glycero-3-phosphocholine (DPPC), 1,2-dipalmitoyl-sn-glycero-3-phosphoethanolamine (DPPE) 1,2-dioleoyl-sn-glycero-3-phosphocholine (DOPC), 1,2-dioleoyl-sn-glycero-3-phosphoethanolamine (DOPE) and brain L-α-phosphatidylinositol-4,5-bisphosphate (ammonium salt) PI(4,5)P_2_ were purchased as powder from Avanti Polar Lipids (purity >99%, Alabaster, AL, USA). Lipid solutions were prepared in chloroform stabilised with ethanol (purity 99.8%; Sigma-Aldrich, St. Louis, MO, USA). Ultra-pure water was generated by passing deionised water through a Milli-Q unit (total organic content = 4 ppb; resistivity = 18 mΩ·cm, Milli-Q, Merck KGaA, Darmstadt, Germany). D_2_O (99.9% of isotopical purity) was purchased from Sigma-Aldrich (St. Louis, MO, USA) and used as received. Experiments were performed in HKM buffer (25 mM HEPES pH 7.2, 125 mM potassium acetate, 5 mM magnesium acetate, 1 mM dithiothreitol, DTT) or HEPES-NaCl buffer (5 mM HEPES pH 7.0, 150 mM sodium chloride). HEPES (in solution, 1 M in H_2_O, and powder, purity 99.5%), potassium acetate (purity≥99.0%), magnesium acetate (purity≥99.0%), DTT (purity≥99.0%), sodium chloride (purity≥99.0%) and magnesium chloride (purity≥98.0%) were purchased from Sigma Aldrich (St. Louis, MO, USA).

### Protein expression and purification

Protein expression and purification protocols can be found elsewhere ^6,16^. Hydrogenous protein constructs were expressed and purified at the Cambridge Institute for Medical Research, while the deuterated CALM was produced in the ILL D-Lab (deuteration proposal DL-03-260), and subsequently purified in Cambridge.

### Langmuir monolayer deposition

A solution of DPPC:DPPE:PtdIns(4,5)P_2_ (7:2:1 molar ratio) in chloroform at 0.1 mg/mL was used for all the experiments with lipid monolayers. A Langmuir trough (Kibron, Helsinki, Finland) with a maximum area of 166.4 cm^2^ equipped with two coupled symmetric sliding barriers was used to measure the surface pressure (Π) – area (A) isotherms of lipid monolayers. The variation of surface pressure was recorded with a force tensiometer using a Wilhelmy plate made of filter paper (Whatman CHR1 chromatography paper). The subphase was made of HKM buffer (120 mL) and the temperature was maintained at 21.0 ± 0.5 °C for all the experiments.

### Neutron reflectometry on mono- and bilayer model membranes

NR experiments were performed on the reflectometers FIGARO^69^ and D17 at the ILL. Two different configurations of angles of incidence were employed: 0.62° and 3.8° (for Langmuir monolayers), and 0.8° and 3.2° (for solid supported bilayers) with a wavelength resolution of 7% dλ/λ on FIGARO and varying from 1 to 20% on D17, leading to a momentum transfer, q_z_=(4π/λ)sinθ, range from 0.001 to 0.25 A^-1^, the higher q_z_ value being determined by the experimental background. Lipid monolayers were prepared in a Langmuir trough, and SLBs by vesicle fusion onto highly polished hydrophilic silicon substrates with a 8×5cm^2^ surface, and after their characterisation, protein was injected in the bulk phase to a final concentration of 5 µM that corresponds to the protein K_D_^3^. For Langmuir monolayers, three buffered isotopic solvent contrasts were employed, characterised by different scattering length densities (SLD): *v/v* 100% D_2_O (SLD = 6.36·10^-6^ Å^-2^), 60% D_2_O (SLD = 3.59·10^-6^ Å^-2^) and 8.1% D_2_O *v/v* (SLD = 0), the last one denominated air contrast matched water (ACMW), since its SLD is equal to that of the air. For SLBs, the contrasts employed were 100% H_2_O (SLD = -0.56·10^-6^ Å^-2^), 38% D_2_O (SLD = 2.07·10^-6^ Å^-2^) and 62% D_2_O v/v (SLD = 3.7·10^-6^ Å^-2^) and 100%D_2_O. (See ***Supplementary Information***)

The NR data were reduced and normalised using COSMOS^70^. Subsequent data analysis was performed using AuroreNR^71^ and Motofit^72^ pieces of software, by minimising the difference between the experimental data points and the calculated reflectivity profile, which was obtained employing a multi-layer slab model (see **SI** for specific details). All the fixed and fitting parameters are tabulated in ***Tables S1***, while the fitting ones (thickness of the slabs, solvent fraction, roughness) are listed in ***Tables S2*** *– **S4***. The software SLDMOL^73,74^ was employed to refine the NR data analysis, by employing atomistic protein models that can be differently oriented with respect to the lipid monolayer, allowing the determination of the protein orientation that can best fit the experimental NR data.

### Grazing Incidence X-ray diffraction (GIXD)

For the GIXD experiments, Langmuir monolayers were irradiated with 8 keV X-ray beam energy and at an incidence angle of θ = 0.1233°, 80% below the critical angle of pure water. GIXD 2D contour profiles of the scattered intensity were acquired using a double linear detector (Mythen 2K) mounted behind a vertically oriented Sollers collimator with an in-plane angular resolution of 1.4 mrad. Diffracted intensities were detected as a function of X-ray momentum transfer component perpendicular, q_z_, and parallel to the air/water interface, q_xy_ = (4π⁄λ) sin 2θ_*xy*_/2, which further gives the repeat distance d = 2π/q_xy_ ,where 2θ_xy_ is the angle between the incident and diffracted beam projected on the air/water interface. The in-plane component reports the lateral crystalline order in the acyl chains of the phospholipid molecules, whereas the out-of-plane component is used to determine the acyl chain tilting angle and coherence length.

GIXD peaks were obtained by the integration of the 2D profiles along q_z_ to obtain the so-called Bragg peaks using in-house scripts developed at ESRF ID10. These data were fitted using Voigt functions to obtain unit cell parameters. The in-plane coherence length (L_xy_) along the crystallographic direction was determined using the Scherrer formula: L_xy_ = (0.9 x 2π)/FWHM_xy_, where FWHM_xy_ is the full width half maxima calculated from the Voigt fitting of the q_xy_ spectrum, respectively.

### Atomic Force Microscopy

Atomic Force Microscopy (AFM). All images were acquired using a multimode AFM with a Nanoscope V controller (Bruker). The AFM was operated in Peak Force Quantitative Nanomechanics (QNM) mode in buffer, at 21.0 ± 0.5 °C. A silicon tip (PNP-DB, NanoAndMore GmbH, Germany), with a spring constant of 0.48 N/m and a resonant frequency of 300 kHz, was used for scanning. Imaging was conducted at a scan rate of 1 Hz with a resolution of 512 × 512 pixels using Nanoscope Software. WSxM software was utilised for the analysis of all AFM images.The AFM tip was calibrated using a standard substrate of sapphire and HOPG in air, followed by calibration in water on the mica surface.

### Quartz-crystal microbalance with dissipation monitoring (QCM-d) measurements

QCM-d measurements were performed using a Q-Sense E4 instrument (Biolin Scientific, Sweden). Frequency and dissipation responses were analysed with QSense Dfind software (Biolin Scientific, Sweden). Changes in resonance frequency (Δf) were recorded for the third, fifth, seventh, ninth, and eleventh overtones. The data presented correspond to the fifth overtone, with variations in Δf and ΔD between overtones being larger than the signal noise, *i.e.*, the data for the different overtones do not overlap, which allows discussing potential viscoelastic effects. Samples were degassed before being introduced into the QCM flow module.

### Sample preparation for NMR experiments

Liposomes were obtained by mixing DOPC and PtdIns(4,5)P_2_ phospholipids at the desired ratio in chloroform (DOPC/PtdIns(4,5)P_2_ molar ratio of 96/4). Chloroform was removed at room temperature with a gentle N_2_ flow followed by drying in a desiccator connected to a vacuum pump (Residual organic solvent was removed by pumping for at least 2 h with a mechanical vacuum pump). D_2_O buffer composed of HEPES buffer (20 mM Hepes, 170 mM NaCl, 2 mM EDTA, pH 7.2) was used to hydrate the lipid film up to a final concentration of 0.50 mg/mL. DOPC: PtdIns(4,5)P_2_ liposomes samples were left overnight (∼12 hours) at room temperature with end-over-end rotation. The liposomes were then packed inside a 4.0 mm zirconia solid-state NMR rotor, through centrifugation using a custom-made 3D-printed centrifugal device for solid-state NMR rotors (STL files available in ***Supporting Information***), based on previously published designs ^75^.

The MLVs used with CALM protein where prepared following the methodology described above. For protein incubation, the hydrated liposomes were placed in a thermostatic water bath at 25 °C and CALM protein buffer solution (HEPES buffer 20 mM Hepes, 170 mM NaCl, 2 mM EDTA, pH 7.2, prepared in H_2_O) was added to the liposomes, yielding a final ratio of PtdIns(4,5)P_2_:CALM of 5:1. The liposomes and protein where kept at 25 °C under constant stirring for a period of 24 h to maximize CALM binding. The incubated liposomes were then centrifuged and packed inside a 4.0 mm zirconia solid-state NMR rotor.

The solid-state NMR experiments were performed immediately after sample preparation to prevent sample aging effects.

### Solid-state NMR Experiments

All the NMR measurements were performed on a Bruker Avance III 700 MHz narrow-bore spectrometer operating at a B_0_ field of 16.4T with ^1^H, ^13^C and ^31^P Larmor frequencies of 700.130, 176.048 and 283.419 MHz, respectively. The experiments were performed on a double-resonance 4.0 mm Bruker MAS probe. The samples were packed into ZrO_2_ rotors with Kel-F caps. Spinning rates between 2 and 10 kHz were employed to record all the spectra (see figure captions for details). ^1^H and ^13^C are quoted in ppm from TMS (0 ppm) and *α*-glycine (secondary reference, C=O at 176.50 ppm). ^31^P CSs were referenced with respect to the external secondary reference phosphoric acid (H_3_PO_4_) at 0.0 ppm. All experiments were performed with temperature control set at 285 K.

^1^H single-pulse excitation under magic-angle spinning (MAS) NMR spectra were recorded at a spinning rate of 8.0 kHz using a pulse width of 4.0 µs (90° flip angle), corresponding to a radio frequency (RF) field strength of 62.5 kHz. A recycle delay (RD) of 2 second was found to be sufficient and used for all samples.

^13^C single-pulse excitation MAS experiments used the same RD, MAS frequency and RF field strength used for the ^1^H single-pulse experiments. The ^13^C cross-polarisation (CP) MAS spectra were recorded under the following conditions: ^1^H 90° pulse set to 4.0 µs corresponding to a radio frequency (RF) field strength of 62.5 kHz; the CP step was performed with a contact time (CT) of 6500 ms using a 70–100% RAMP shape at the ^1^H channel and using a 50 - 75 kHz square shape pulse on the ^13^C channel; RD was 2 s. During the acquisition, a SPINAL-64 decoupling scheme was employed using a pulse length for the basic decoupling units of 6.250 µs at an RF field strength of ∼73.33 kHz.

The ^13^C-{^1^H} CP HETCOR spectra were recorded using the CP conditions described above. 410 t1 points with 608 scans were recorded along the indirect dimension. Quadrature detection in t1 was achieved using States-TPPI.

^31^P single-pulse excitation magic angle spinning (MAS) NMR spectra were recorded at a spinning rate from 2-8 kHz using a pulse width of 6.30 µs (90° flip pulse) corresponding to a radio frequency (rf) field strength of ∼ 39.7 kHz. A recycle delay of 5 s was found to be sufficient and used for all samples.

## Acknowledgements

The authors thank both the Institut Laue-Langevin (DOI:ILL-DATA.8-02-829) and the European Synchrotron Radiation Facility ESRF (DOI:10.15151/ESRF-ES-883953841) for the allocation of beamtime and the Partnership for Soft Condensed Matter (PSCM) for the lab support. A.M. acknowledges the financial support from MCIN/AEI/10.13039/501100011033 under grant PID2021-129054NA-I00, and from the Department of Education of the Basque Government under grant PIBA_2023_1_0054 and from the IKUR Strategy under the collaboration agreement between Ikerbasque Foundation and Materials Physics Center. E.G. acknowledge the financial support MCIN/AEI/10.13039/501100011033 and UCM under grants PID2019-106557GB-C21 and PR12/24-31566 (Ayudas para la financiación de proyectos de investigación UCM 2023). NZ and DJO acknowledge the support of the Wellcome Trust grant to DO (207455/Z/17/Z). This work was also developed within the scope of project CICECO-Aveiro Institute of Materials, UIDB/50011/ 2020, UIDP/50011/2020 & LA/P/0006/2020, financed by national funds through the FCT/MEC (PIDDAC). The NMR spectrometers are part of the National NMR Network (PTNMR) and are partially supported by Infrastructure Project 022161 (cofinanced by FEDER through COMPETE 2020, POCI and PORL and FCT through PIDDAC). FCT is also acknowledged by D. P. for a Ph.D. Studentship UI/BD/151048/2021 (DOI: 10.54499/UI/BD/151048/2021). I.M.-M. acknowledges the EMBO organisation for the EMBO Fellowship 8740.

