## Supplemental material (text, figures, tables) for "Untangling structural molecular details of the endocytic adaptor protein CALM upon binding with phosphatidylinositol 4,5-bisphosphate-containing model membranes"

### Experimental Details

##### *Surface pressure (Π) - area (A) isotherm of DPPC:DPPE:PtdIns(4,5)$P_2$ monolayer*

The surface pressure (Π) - area (A) per molecule compression isotherm of the DPPC:DPPE: PtdIns(4,5)$P_2$ (molecular structures are shown in **Figure S1**) monolayer was recorded, at 21°C, and plotted in **Figure 1**. The corresponding compressional elastic moduli ($C_s^{-1}$) plotted also in **Figure 1** was calculated according to **Equation S1**.

$$C_s{}^{-1} = -\mathrm{A} \cdot \left(\frac{\partial \Pi}{\partial \mathrm{A}}\right) \qquad \textbf{S1}$$

The compression isotherm allowed us to explore a large region of the lipid interfacial phase diagram, providing insightful knowledge about the monolayer phase behaviour.

##### *Chemical Shift Anisotropy Tensor Convention*

The NMR chemical shift exhibits variations in its magnitude depending on the arrangement of atoms around the nucleus. This variation, known as chemical shift anisotropy (CSA), is influenced by the orientation of the molecule concerning an external magnetic field. Mathematically, CSA is characterized by a second-order tensor, typically represented as a 3 x 3 matrix (**Eq. S2**). It is common practice to express this tensor in a coordinate system where all non-diagonal elements are zero. In this system, the chemical shift tensor is fully defined by three diagonal elements, termed principal components ($\boldsymbol{\delta_{XX}}$, $\boldsymbol{\delta_{YY}}$, $\boldsymbol{\delta_{ZZ}}$), and the orientation of their corresponding principal axes, which are described by three Euler angles.

$$\widehat{\boldsymbol{\delta}} = \begin{bmatrix} \boldsymbol{\delta_{XX}} & \boldsymbol{\delta_{YX}} & \boldsymbol{\delta_{ZX}} \\ \boldsymbol{\delta_{XY}} & \boldsymbol{\delta_{YY}} & \boldsymbol{\delta_{ZY}} \\ \boldsymbol{\delta_{XZ}} & \boldsymbol{\delta_{YZ}} & \boldsymbol{\delta_{ZZ}} \end{bmatrix} \qquad \textbf{S2}$$

In this work the Haeberlen convention is adopted to express the CSA tensor [1,2]. In Haeberlen convention, the principal values of the CSA are ordered according to the distance from the isotropic value ($\boldsymbol{\delta_{iso}}$) as follows:

$$|\boldsymbol{\delta_{ZZ}} - \boldsymbol{\delta_{iso}}| \geq |\boldsymbol{\delta_{XX}} - \boldsymbol{\delta_{iso}}| \geq |\boldsymbol{\delta_{YY}} - \boldsymbol{\delta_{iso}}| \qquad \textbf{S3}$$

Where $\boldsymbol{\delta_{iso}}$ is defined as:

$$\boldsymbol{\delta_{iso} = (\delta_{XX} + \delta_{YY} + \delta_{ZZ})/3} \quad \textbf{S4}$$

Reduced anisotropy ($\boldsymbol{\delta}$) is the largest separation from the isotropic value and is defined as:

$$\boldsymbol{\delta = \delta_{ZZ} - \delta_{iso}} \quad \textbf{S5}$$

Asymmetry parameter ($\boldsymbol{\eta}$) describes how much the NMR line shape deviates from an axially symmetric tensor and is described as:

$$\boldsymbol{\eta = (\delta_{YY} - \delta_{XX})/\delta} \quad \textbf{S6}$$

In the case of an axially symmetric tensor $\boldsymbol{\delta_{YY} - \delta_{XX}}$ will be zero and hence $\boldsymbol{\eta}$ = 0.

Additional information regarding other CSA conventions and how to convert values between them can be found in *Chemical Shift Tensor Conventions* article in the webpage of Universität Tübingen.

#### *Solid-supported lipid bilayers (SLBs) preparation*

SLBs were prepared by PtdIns(4,5)$P_2$-enriched liposome adsorption and fusion into a solid substrate. Small unilamellar vesicles (SUVs) were prepared by dissolving DOPC:DOPE: PtdIns(4,5)$P_2$ (7:2:1 molar ratio) in chloroform, dried under gentle Argon flow and placed in vacuum overnight to ensure evaporation of all solvent. The resulting lipid films were rehydrated at room temperature in HKM or HEPES buffer solutions up to 0.1 mg·$mL^{-1}$. The suspension was tip sonicated for 5 min at pulses of 1 s on/off to produce a visually clear dispersion of SUVs and then extruded at room temperature through 100 nm pore-size Millipore polycarbonate membranes. The suspension was introduced in the flow cell at 0.1 mL·$min^{-1}$ and the typical signal corresponding to the vesicle fusion and lipid bilayer formation was followed. Osmotic shock was exploited to force vesicle rupture and bilayer formation, by introducing in the flow cell a solution 500 mM of sodium chloride at 0.1 mL·$min^{-1}$. After the formation of the lipid bilayer, buffer solution was flowed through the system to remove partially or weakly bound unilamellar lipid vesicles. Protein solution (9.6 µM) was introduced in the flow cell at 0.1 mL·$min^{-1}$. After 5 hours incubation, buffer was flowed at 0.1 mL·$min^{-1}$ to remove unbound and physisorbed protein molecules. The system was let equilibrate overnight and finally buffer was flowed at 0.1 mL·min$^{-1}$ until a stable baseline was reached.

##### *Further description of Neutron reflectometry (NR) experiments*

NR experiments were performed on the reflectometers FIGARO[3] and D17[4,5] at the ILL. Two different angles of incidence were employed: 0.62° and 3.8° for Langmuir monolayers, and 0.8° and 3.2° (or 3°) for solid supported lipid bilayers in FIGARO (or D17). The wavelength resolution used was 7% dλ/λ on FIGARO and varied from 1 to 20% on D17. The momentum transfer, $q_z=(4\pi/\lambda)\sin\theta$, range was from 0.001 to 0.25 A$^{-1}$.

**Lipid monolayers** were prepared in a Langmuir trough, filled with HKM buffer solution, and after their characterisation, protein was injected in the bulk phase to a final concentration of 5 µM that corresponds to the protein $K_D$[6]. Three buffered isotopic solvent contrasts were employed, characteri by different scattering length density (SLD): 100% $D_2O$ (SLD=6.36·10$^{-6}$ Å$^{-2}$), 60% $D_2O$ *v/v* (SLD=3.59·10$^{-6}$ Å$^{-2}$) and 8.1% $D_2O$ *v/v* (SLD=0), denominated air contrast matched water (ACMW), since its SLD is equal to that of the air.

**NR experiments on solid-supported lipid bilayers**, solid/liquid flow cells available at the ILL with polished silicon crystals (111) with a surface area of 5×8 cm$^2$ were used. The silicon crystals were carefully cleaned with organic solvents (chloroform, acetone, ethanol) and made hydrophilic by 2 minutes plasma cleaning by using a PDC-002 (230V) (Harrick Plasma, New York, US). Substrate surface was characteri in two different isotopic solvent contrasts, (100% $H_2O$ and 100% $D_2O$), before bilayer deposition. Small unilamellar vesicles (SUVs) were prepared by dissolving DOPC:DOPE: PtdIns(4,5)$P_2$ (7:2:1 molar ratio) in chloroform, dried under gentle Argon flow and placed in vacuum overnight to ensure evaporation of all solvent. The resulting lipid films were rehydrated at room temperature in HEPES-NaCl buffer with 2 mM magnesium chloride up to 1 mg·mL$^{-1}$. Immediately before use for lipid bilayer formation, the suspension was tip sonicated for 5 min at pulses of 1 s on/off to produce a visually clear dispersion of SUVs. Bilayers were formed through vesicle fusion, aided by osmotic shock (with pure water), and then characteri in three different buffered isotopic solvent contrasts: 100% $H_2O$ (SLD=-0.56·10$^{-6}$ Å$^{-2}$), 38% $D_2O$ *v/v* (called silicon contrast matched water, SiMW, SLD=2.07·10$^{-6}$ Å$^{-2}$) and 100% $D_2O$. Then, the protein was injected in the flow cell, up to a 20 µM concentration. Unbound and physisorbed protein molecules were washed away by a buffer solution washing step, and the bilayer+protein system was characterised in three buffered isotopic solvent contrasts: 100% $H_2O$, 38% $D_2O$ and

100% $D_2O$ *v/v* for experiments with hydrogenous CALM, and 100% $H_2O$, 38% $D_2O$ and 62% $D_2O$ *v/v* for experiments with deuterated CALM. HEPES 5mM, NaCl 150mM (pH=7) buffer solution was used for NR bilayer experiments. The NR data were reduced and normali using COSMOS[7].

**NR Data analysis** was performed using AuroreNR[8] and Motofit[9] software, by minimizing the difference between the experimental data points and the calculated reflectivity profile, which was obtained employing a multi-layer slab model. The latter is made of layers characteri by an in-plane averaged SLD, with value depending on the volume fraction (*f*) of each component, leading to $SLD_{model} = f_{solvent} \cdot SLD_{solvent} + f_{lipid} \cdot SLD_{lipid} + f_{protein} \cdot SLD_{protein}$; where $f_{solvent} + f_{lipid} + f_{protein} = 1$. Data analysis was performed using constraints between layer parameters, *i.e.*, taking into account that lipid headgroups and tails must have the same area per molecule. To this purpose, the fraction of water in the headgroups-layer ($f_{wh}$) was calculated following $f_{\mathrm{wh}} = 1 - \frac{\mathrm{t_t} \cdot \mathrm{V_h}}{\mathrm{t_h} \cdot \mathrm{V_t}}$, where $t_t$ and $t_h$ are the thicknesses of tails- and headgroups-layer, respectively, and $V_t$ and $V_h$ are the molecular volumes of tails and headgroups, respectively. All the fixed parameters are tabulated in **Table S1**. The roughness of the interfaces was also fitted; in the case of purely lipid molecules containing layer, the roughness of the different interfaces was assumed to be identical.

The variation of the volume fraction, Φ(z), for each component with the distance to the interface, *z*, was calculated using the model composed of N-layers of varying volume fraction ($f_i$) and thickness ($t_i$) modulated by a roughness parameter ($\sigma_i$) which describes the interfacial mixing of the layers, following $\Phi(\mathrm{z}) = \sum_{i=0}^{N} \frac{f_i - f_{i-1}}{2} \left(1 + erf\left(\frac{z - t_i}{\sigma_i \sqrt{2}}\right)\right)$.

In a final level of refinement, SLDMOL program [10,11] was employed to refine the NR data analysis, by employing atomistic protein models that can be differently oriented with respect to the lipid monolayer. Thus, it allows the determination of the protein orientation that can best fit the experimental NR data. SLDMOL software is a freely distributed open-source program written in python, and is available on the online SASSIE package platform (https://sassie-web.chem.utk.edu/sassie2) [10,12].

### Tables

**Table S1.** Fixed parameters for mono- and bilayers used for the NR data analysis[13,14]. The exchange of protons in the contrasts with different $D_2O$ content was taken into account. In the case of lipid monolayer, the volume contraction for palmitoyl tails was taken into account [15].

| **Fixed Parameters** | DPPC:DPPE:PtdIns(4,5)$P_2$ (7:2:1) monolayer | DOPC:DOPE:PtdIns(4,5)$P_2$ (7:2:1) bilayer |
|---|---|---|
| $V_h$ (Å$^3$) | 320.9 | 320.9 |
| $SLD_h$ ($10^{-6}$ Å$^{-2}$) | 2.17 | 2.17 |
| $V_t$ (Å$^3$) | 732.3 | 987 |
| $SLD_t$ ($10^{-6}$ Å$^{-2}$) | -0.42 | -0.2* |

**Table S2.** Fitting parameters of lipid monolayers at different surface pressures (15 and 25 mN·m$^{-1}$) and lipid bilayers with $CALM_{wt}$. The $\chi^2$ of all fits reported here are below 10.

| **Monolayer** | $\Pi_0$=15mN·m$^{-1}$ | | | | $\Pi_0$=25mN·m$^{-1}$ | | | |
|---|---|---|---|---|---|---|---|---|
| | t (Å) | $f_w$ (%) | r (Å) | APM (Å$^2$) | t (Å) | $f_w$ (%) | r (Å) | APM (Å$^2$) |
| Tails-layer | 12±1 | | 4±1 | 63±5 | 15±1 | | 5±1 | 49±3 |
| Headgroups-layer | 8±1 | 35±1 | 4±1 | 59±8 | 9±1 | 24±1 | 5±1 | 47±5 |

**Table S3.** Thickness and volume fraction of water ($f_w$) and protein (fitting parameters), for $CALM_{wt}$ and $CALM_{\Delta AH0}$ interacting with monolayers at $\Pi_0$=15mN·m$^{-1}$. The roughness (r) of the interfaces and the area per molecule (APM) of lipids are also reported. The total protein coverage obtained with SLDMOL is 12% and 15% (*v/v*) for $CALM_{wt}$ and $CALM_{\Delta AH0}$, respectively.

| | t (Å) | $f_w$ (%) | $CALM_{wt}$ % (*v/v*) | r (Å) | APM (Å$^2$) | t (Å) | $f_w$ (%) | $CALM_{\Delta AH0}$ % (*v/v*) | r (Å) | APM (Å$^2$) |
|---|---|---|---|---|---|---|---|---|---|---|
| Tails-layer | 12±1 | | 0 | 5±1 | 59±5 | 12±1 | | 0 | 3±1 | 59±5 |
| Headgroups-layer | 7±1 | 11±1 | 13±1 | 5±1 | 56±9 | 8±1 | 28±1 | 5±1 | 3±1 | 58±9 |
| CALM-layers | 17±1 | 88±1 | 12±1 | 3±1 | | 20±1 | 84±1 | 16±1 | 8±1 | |
| | 15±1 | 92±1 | 8±1 | 3±1 | | 15±1 | 90±1 | 10±1 | 8±1 | |
| | 18±1 | 93±1 | 7±1 | 3±1 | | 20±1 | 92±1 | 8±1 | 8±1 | |

**Table S4.** Thickness and volume fraction of water ($f_w$) and protein, for h- and d- $CALM_{wt}$ interacting with bilayers.

| **Bilayer** | t (Å) | $f_w$ (%) | r (Å) | APM (Å$^2$) | t (Å) | $f_w$ (%) | $CALM_{wt}$ % (*v/v*) | r (Å) | APM (Å$^2$) |
|---|---|---|---|---|---|---|---|---|---|
| Water-layer | 6±1 | 100 | 4±1 | | 4±1 | 100 | 0 | 4±1 | |
| Headgroups-layer | 8±1 | 39±1 | 4±1 | 64±9 | 9±1 | 48±1 | 0 | 4±1 | 71±9 |
| Tails-layer | 16±1 | 5±1 | 4±1 | 63±5 | 15±1 | 6±1 | 0 | 4±1 | 70±5 |
| Tails-layer | 16±1 | 11±1 | 4±1 | 68±5 | 16±1 | 16±1 | 0 | 4±1 | 74±6 |
| Headgroups-layer | 7±1 | 50±1 | 4±1 | 71±12 | 8±1 | 48±1 | 3±1 | 4±1 | 78±11 |
| d- CALM-layer | | | | | 56±1 | 96±1 | 4±1 | 10±1 | |

| **Bilayer** | t (Å) | $f_w$ (%) | r (Å) | APM (Å$^2$) | t (Å) | $f_w$ (%) | $CALM_{wt}$ % (*v/v*) | r (Å) | APM (Å$^2$) |
|---|---|---|---|---|---|---|---|---|---|
| Water-layer | 4±1 | 100 | 3±1 | | 5±1 | 100 | 0 | 3±1 | |
| Headgroups-layer | 8±1 | 44±1 | 3±1 | 70±10 | 8±1 | 43±1 | 0 | 3±1 | 70±10 |
| Tails-layer | 30±1 | 4±1 | 3±1 | 69±3 | 14±1 | 3±1 | 0 | 3±1 | 71±6 |
| Tails-layer | | | | | 14±1 | 6±1 | 0 | 3±1 | 73±6 |
| Headgroups-layer | 8±1 | 44±1 | 3±1 | 70±10 | 8±1 | 39±1 | 3±1 | 5±1 | 75±11 |
| h-CALM-layer | | | | | 56±1 | 96±1 | 4±1 | 5±1 | |

**Table S5.** GIXD structural parameters obtained for CALM$_{wt}$ interacting with monolayers at $\Pi_0$=15mN·m$^{-1}$.

| **Monolayer** | **Unit cell parameters** | | | **In Plane Bragg Peaks** | | | | |
|---|---|---|---|---|---|---|---|---|
| | **a/b (Å)** ±0.05 Å | **γ ( deg )** ±1.0 $^0$ | **Chain cross-sectional area** ±0.5 (Å$^2$) | **$d_{01}$ / $d_{10}$ (Å)** ±0.05 Å | **$d_{11}$ (Å)** ±0.05 Å | **$L_{01}$/$L_{10}$ (Å)** | **$L_{11}$ (Å)** | **φ** **(**deg**)** ± 0.5 $^0$ |
| **DPPC / DPPE / PtdIns(4,5)P$_2$** | 5.08 | 114 | 24 | 4.63 | 4.26 | 78 ± 2 | 263 ± 5 | 40.2 |
| **DPPC / DPPE / PtdIns(4,5)P$_2$ + CALM** | 4.99<br>4.66 | 114 | 21 | 4.56<br>4.03 | 4.26 | 153 ± 3<br>99 ± 5 | 239 ± 5 | 33.5 |

**Table S6.** $^{31}$P direct excitation ssNMR spectral deconvolution parameters of DOPC/PtdIns(4,5)P$_2$ liposomes in the absence and presence of CALM$_{wt}$ protein. The $^{31}$P spectra is shown in ***Figure S7***.

| **Before CALM$_{wt}$ incubation** | | |
|---|---|---|
| ***Lipid*** | $\delta$ (iso), ppm | L.B., Hz |
| DOPC diester phosphate | -0.89 | 95.35 |
| DOPC diester phosphate (broad) | -0.93 | 452.23 |
| PtdIns(4,5)P$_2$ diester phosphate | 0.34 | 25.37 |
| PtdIns(4,5)P$_2$-P5 | 1.02 | 117.89 |
| PtdIns(4,5)P$_2$-P4 | 2.06 | 120.21 |
| **After CALM$_{wt}$ incubation** | | |
| DOPC diester phosphate | -0.83 | 110.10 |
| DOPC diester phosphate (broad) | -0.80 | 422.77 |
| PtdIns(4,5)P$_2$ diester phosphate | 0.66 | 55.55 |
| PtdIns(4,5)P$_2$-P5 | 1.59 | 175.15 |
| PtdIns(4,5)P$_2$-P4 | 2.71 | 163.62 |

**Table S7.** $^{31}$P longitudinal relaxation $T_1$ time of the phosphate groups of DOPC and PtdIns(4,5)$P_2$ lipids before and after protein incubation.

| | **Before $CALM_{wt}$ incubation** | **After $CALM_{wt}$ incubation** |
|---|---|---|
| ***Lipid*** | $T_1$, s | $T_1$, s |
| DOPC diester phosphate | 0.556 ± 0.012 | 0.577 ± 0.014 |
| PtdIns(4,5)$P_2$ diester phosphate | 0.818 ± 0.104 | 1.007 ± 0.264 |
| PtdIns(4,5)$P_2$-P5 | 0.529 ± 0.085 | 0.334 ± 0.104 |
| PtdIns(4,5)$P_2$-P4 | 0.518 ± 0.174 | 0.371 ± 0.095 |

**Table S8.** $^{31}$P NMR chemical shift anisotropy tensor elements of DOPC and PtdIns(4,5)$P_2$ lipids before and after protein incubation. The $^{31}$P spectra at 4 kHz is shown in ***Figure S9***.

| **Before $CALM_{wt}$ incubation** | | | | |
|---|---|---|---|---|
| ***Lipid*** | $\delta$ (iso), ppm | $\delta$ (CSA), ppm | $\eta$ (CSA), ppm | L.B., Hz |
| DOPC diester phosphate | -0.94 | 30.35 | 0.01 | 193.20 |
| DOPC diester phosphate (broad) | -1.15 | -1.07 | 0.09 | 541.71 |
| PtdIns(4,5)$P_2$ diester phosphate | 0.30 | 0.09 | 0.03 | 101.59 |
| PtdIns(4,5)$P_2$-P5 | 1.02 | -16.5 | 0.04 | 157.98 |
| PtdIns(4,5)$P_2$-P4 | 2.09 | 0.08 | 0.37 | 132.76 |
| **After $CALM_{wt}$ incubation** | | | | |
| ***Lipid*** | $\delta$ (iso), ppm | $\delta$ (CSA), ppm | $\eta$ (CSA), ppm | L.B., Hz |
| DOPC diester phosphate | -0.86 | 28.25 | 0.05 | 186.12 |
| DOPC diester phosphate (broad) | -0.61 | 29.22 | 0.05 | 1052.57 |
| PtdIns(4,5)$P_2$ diester phosphate | 0.64 | -1.24 | 0.09 | 41.62 |
| PtdIns(4,5)$P_2$-P5 | 1.63 | 18.60 | 0.07 | 154.64 |
| PtdIns(4,5)$P_2$-P4 | 2.68 | 8.46 | 0.46 | 207.30 |

### Figures

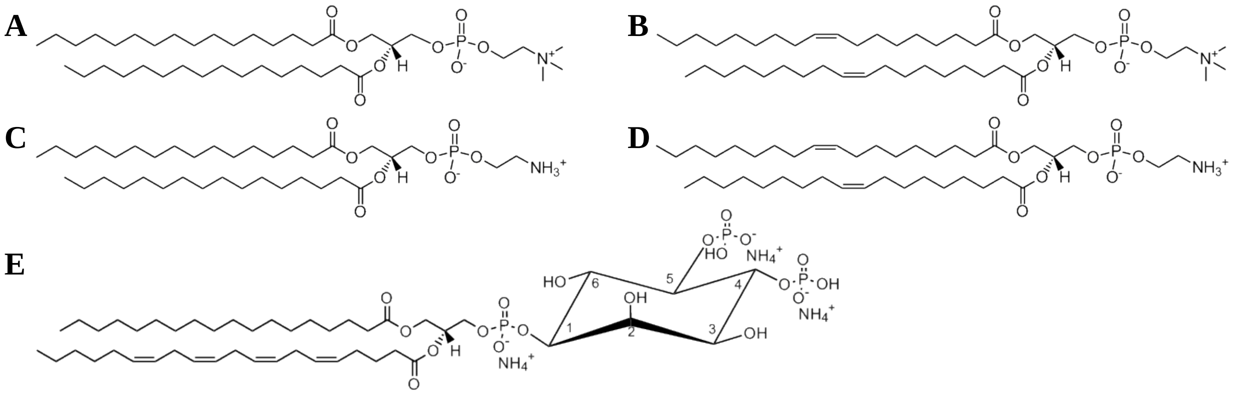

**Figure S1.** Molecular structures of the phospholipids used in this study. **A** DPPC, **B** DOPC, **C** DPPE, **D** DOPE, **E** PtdIns(4,5)$P_2$.

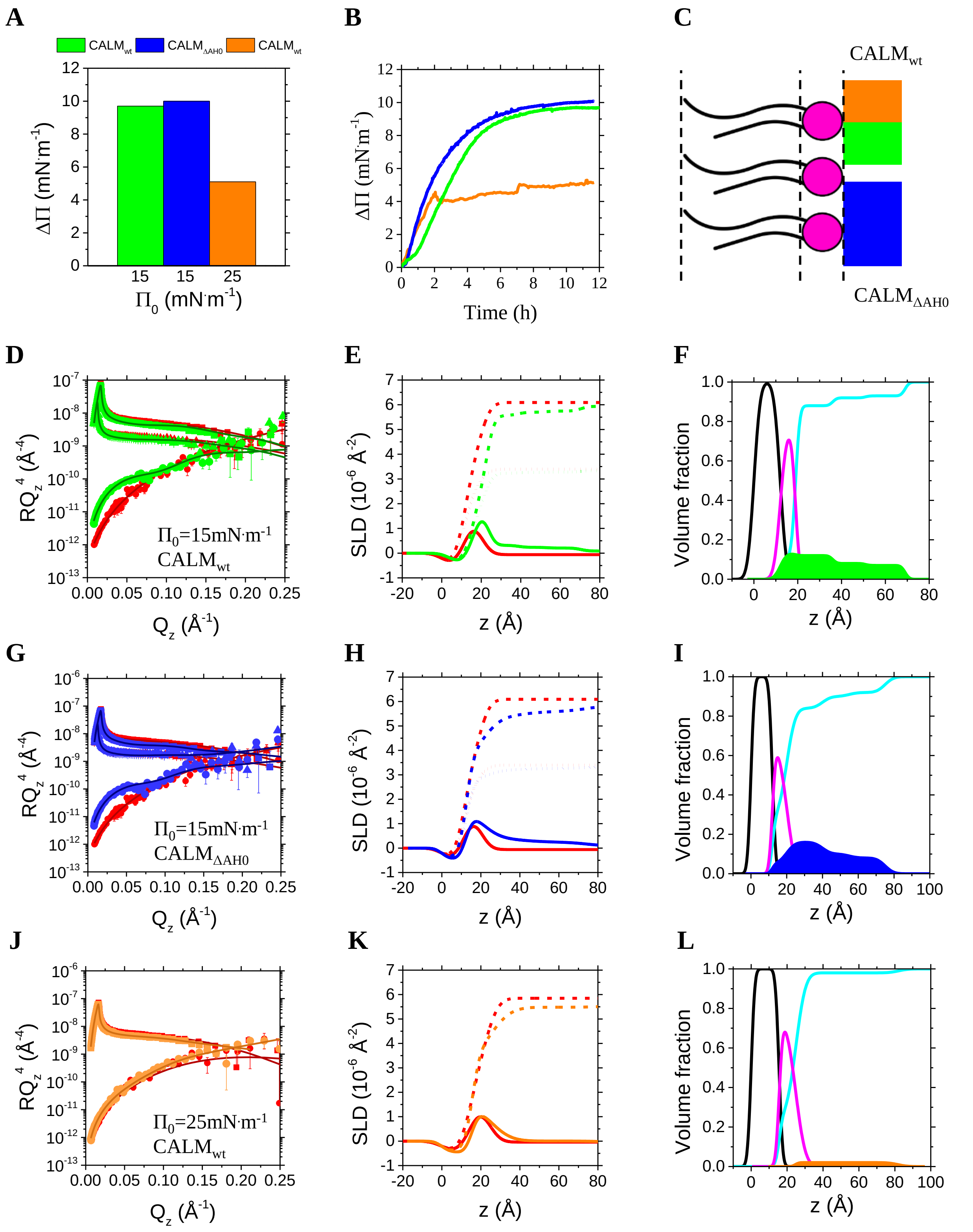


**Figure S2. Experiments with lipid monolayers** (legend below).

Panel **A** shows the increase in surface pressure upon CALM injection in the bulk phase, and panel **B** reports the binding kinetics. Data related to $CALM_{wt}$ injected under a $\Pi_0$=15 mN·m$^{-1}$ and 25 mN·m$^{-1}$ monolayers are depicted in green and orange respectively. Data related to $CALM_{\Delta AH0}$ are depicted in blue. **C** Sketch indicating a lipid monolayer with protein bound underneath. Tails are depicted in black and headgroups in magenta. Experimental (symbols) and simulated (lines) NR profiles of lipid monolayers ($\Pi_0$=15 mN·m$^{-1}$) in the absence (red) and presence of **D** $CALM_{wt}$ (green) and **G** $CALM_{\Delta AH0}$ (blue). Data at three isotopic contrasts have been measured: $D_2O$ (squares), 60% $D_2O$ (triangles) and ACMW (circles). Experimental (symbols) and simulated (lines) neutron reflectivity profiles of lipid monolayers ($\Pi_0$=25 mN·m$^{-1}$) in the absence (red) and presence of **J** $CALM_{wt}$ (orange). Data at two isotopic contrasts have been measured: $D_2O$ (squares) and ACMW (circles). Figures are displayed on an $RQ_z^4$ scale to show the quality of the fits at high $Q_z$ values. SLD profiles corresponding to fits are plotted in **E**, **H** and **K**. Continuous, short dotted, short dashed lines indicate the SLD profiles in ACMW, 60% $D_2O$ and $D_2O$ isotopic contrast, respectively. Red lines in **E** and **H** refer to purely monolayer profiles ($\Pi_0$=15 mN·m$^{-1}$), green and blue lines refer to monolayer+$CALM_{wt}$ and monolayer+$CALM_{\Delta AH0}$ SLD profiles, respectively. Red lines in **K** refer to purely monolayer profiles ($\Pi_0$=25 mN·m$^{-1}$), orange lines refer to monolayer+$CALM_{wt}$ SLD profiles. Volume fraction profiles derived from the fit highlight the distribution of tails (black), heads (magenta), water (cyan) and **F** $CALM_{wt}$ underneath a $\Pi_0$=15 mN·m$^{-1}$ monolayer (green) and **L** a $\Pi_0$=25 mN·m$^{-1}$ monolayer (orange), **I** $CALM_{\Delta AH0}$ underneath a $\Pi_0$=15 mN·m$^{-1}$ monolayer (blue).

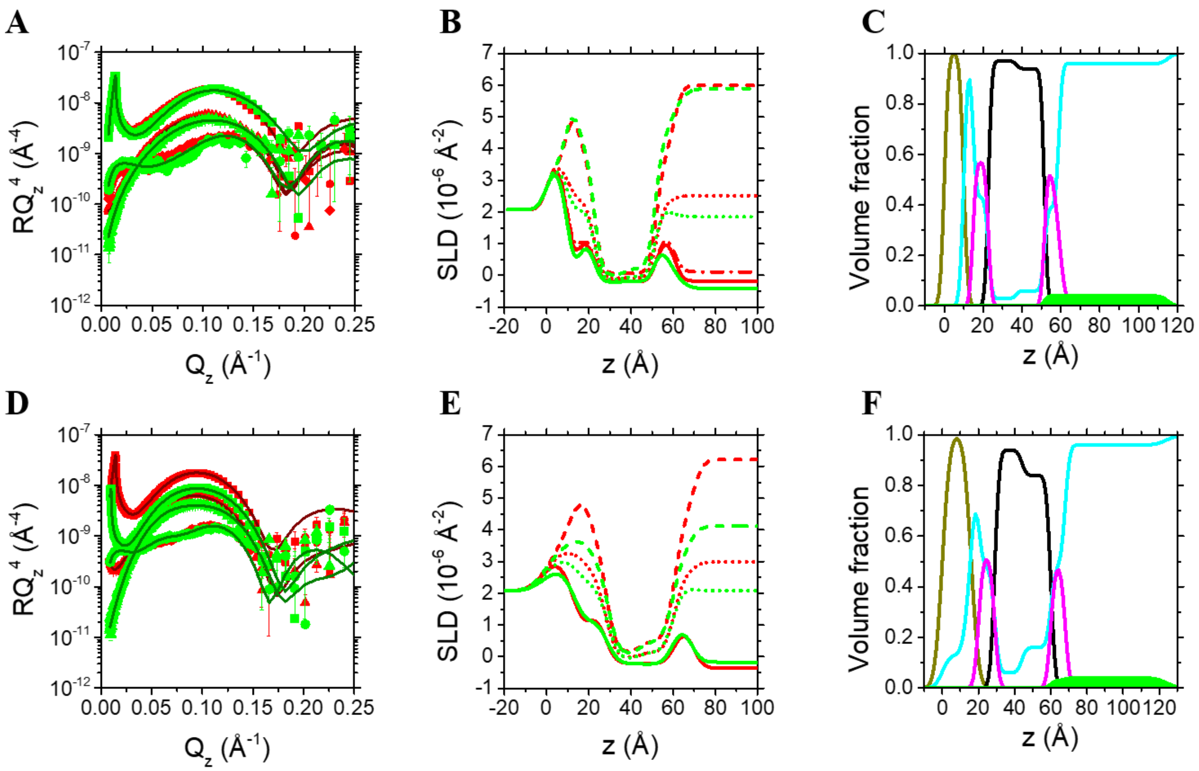


**Figure S3. NR experiments with supported lipid bilayers.** Experimental (symbols) and simulated (lines) neutron reflectivity profiles of solid-supported bilayers in the absence (red) and presence (green) of **A** h-CALM$_{wt}$ and **D** d-CALM$_{wt}$. Data at three isotopic contrasts have been measured for hydrogenous lipid bilayers: $D_2O$ (red squares), $H_2O$ (red circles) and SiMW (red triangles). An additional contrast was measured for the bilayer prior to the injection of h-CALM$_{wt}$, *i.e.*, ACMW, shown as red diamonds. Data at three isotopic contrasts have been measured for bilayers+CALM$_{wt}$: $D_2O$ for h-CALM$_{wt}$ and 62% $D_2O$ for d-CALM$_{wt}$ (green squares), $H_2O$ (green circles) and SiMW (red triangles). Figures are displayed on an $RQ_z^4$ scale to show the quality of the fits at high $Q_z$ values. Scattering length density profiles corresponding to fits are plotted in **B** and **E**. Red continuous, short dotted, short dashed lines indicate the bilayer SLD profile in $H_2O$, SiMW and $D_2O$ isotopic contrast, respectively, and the dotted-dashed line indicated the bilayer SLD in ACMW. Green continuous, short dotted, short dashed lines indicate the bilayer+CALM$_{wt}$ SLD profile in $H_2O$, SiMW and $D_2O$ (60% $D_2O$ for d-CALM$_{wt}$) isotopic contrast, respectively. Volume fraction profiles derived from the fits highlight the distribution of silicon oxide (dark yellow), tails (black), heads (magenta), water (cyan) and **C** h-CALM$_{wt}$ (green), **F** d-CALM$_{wt}$ (green).

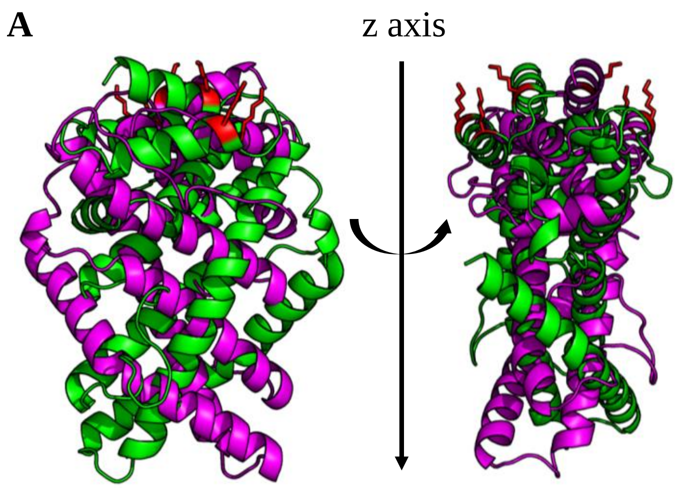


**Figure S4.** Cartoon representation of the two-equivalent orientation of CALM with respect to the z-axis: the orientation α=330°, β=10° is depicted in green, and the one α=150°, β=170° in magenta. The PtdIns(4,5)$P_2$ binding site is red.

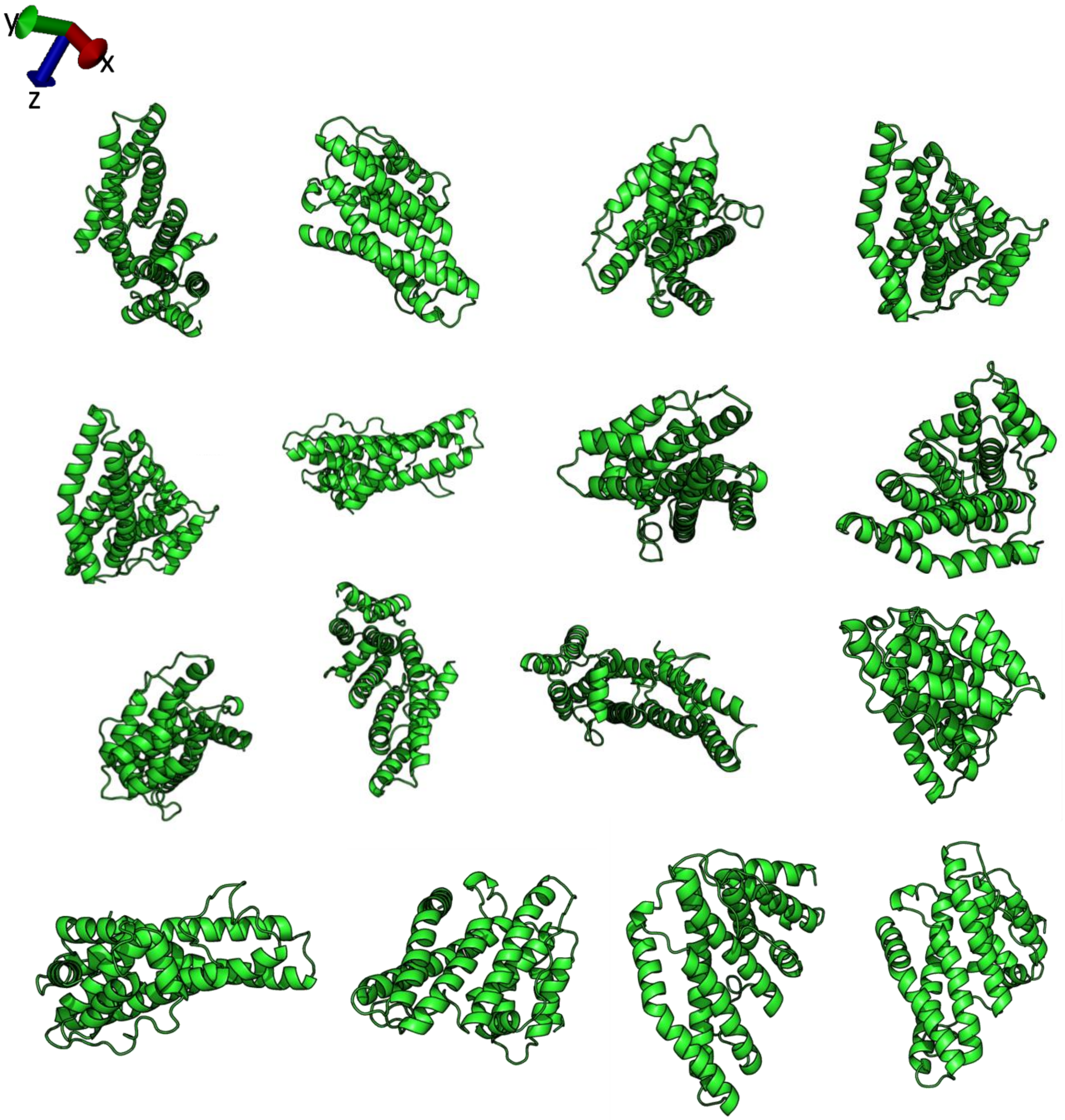


**Figure S5.** Some of the rigid body rotation of CALM (PDB ID: 3ZYK[16]) implemented in the .dcd file.

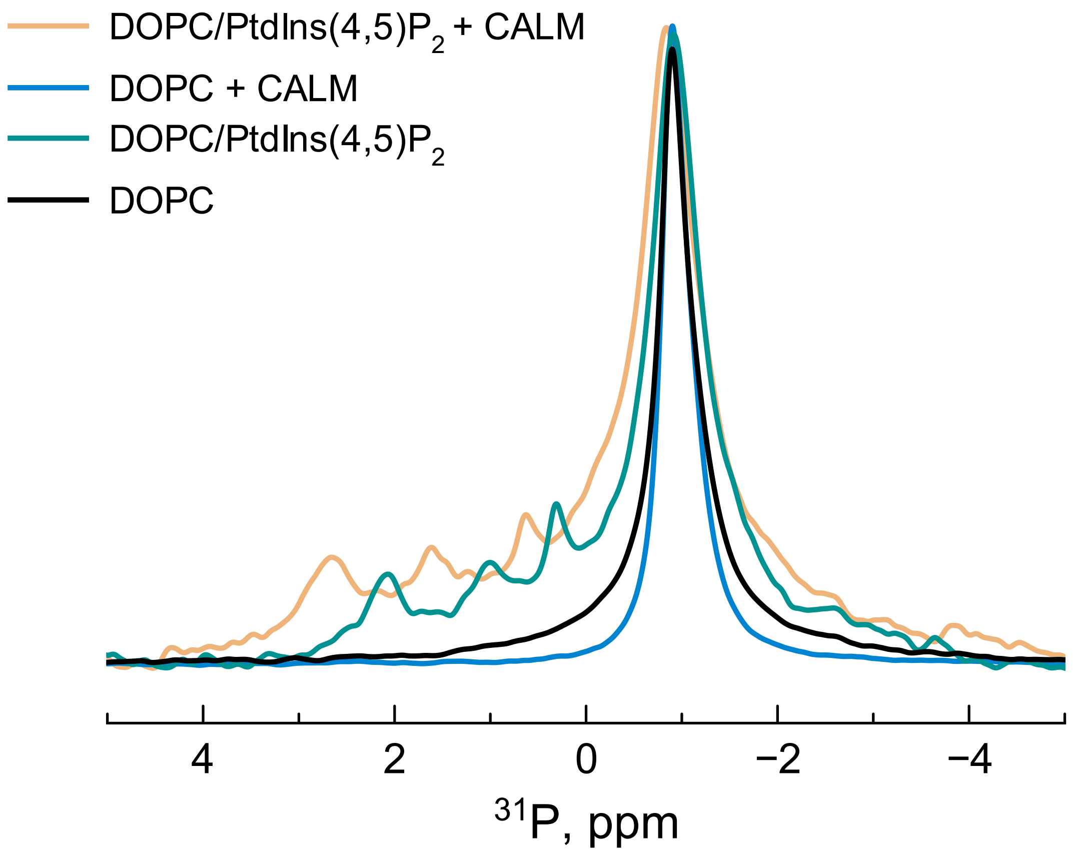


**Figure S6.** $^{31}P$ direct excitation ssNMR spectra of DOPC and DOPC/PtdIns(4,5)$P_2$ liposomes in the presence and absence of $CALM_{wt}$ protein. All the spectra were acquired at 290 K using a MAS speed of 8 kHz.

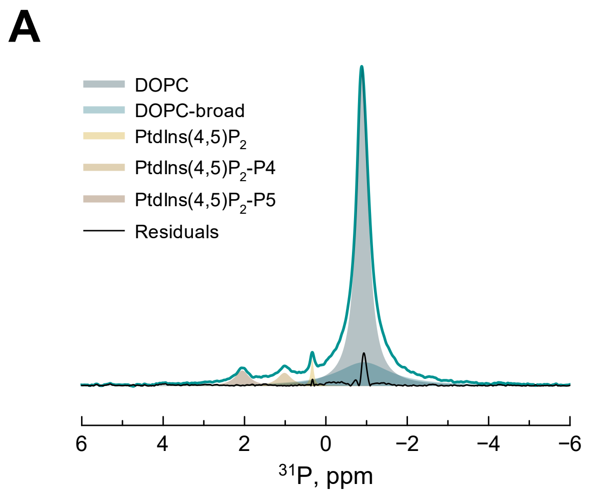


**Figure S7.** $^{31}$P direct excitation ssNMR spectral deconvolution of DOPC/PtdIns(4,5)$P_2$ liposomes in the absence (**A**) and presence of $CALM_{wt}$ protein (**B**). The parameters from the spectra deconvolution are shown in ***Table S6***. All the spectra were acquired at 290 K using a MAS speed of 8 kHz.

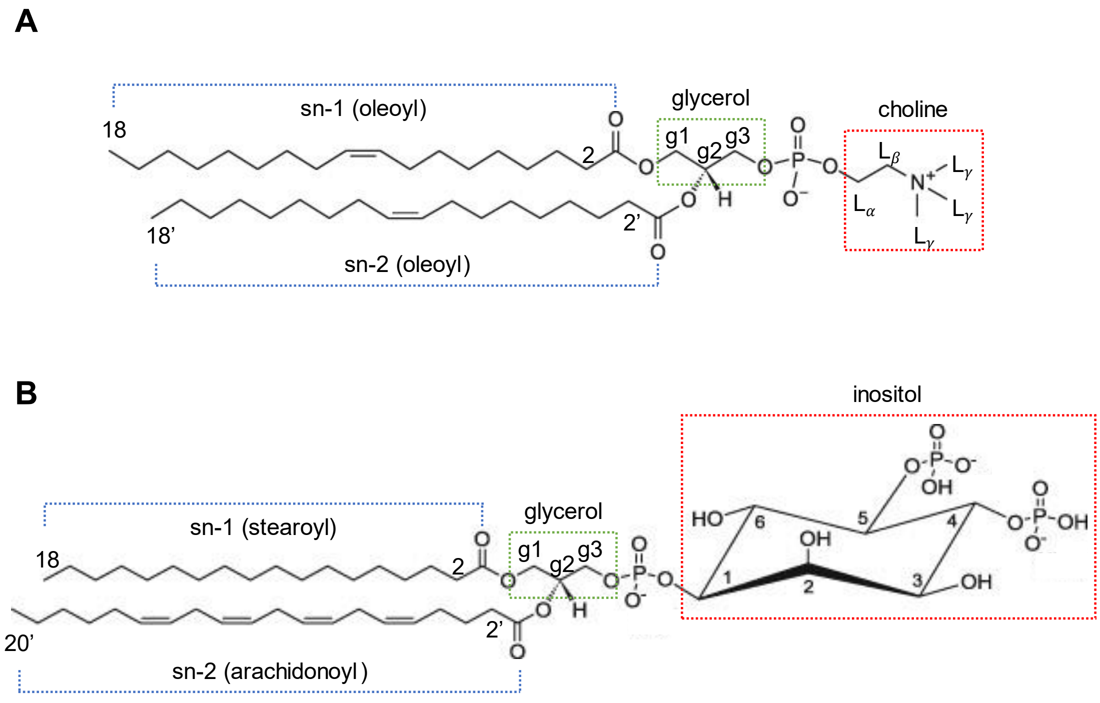


**Figure S8.** Chemical structure of (**A**) DOPC and (**B**) PtdIns(4,5)$P_2$. The chemical group legend was adopted in the assignments of ***Figure 3***.

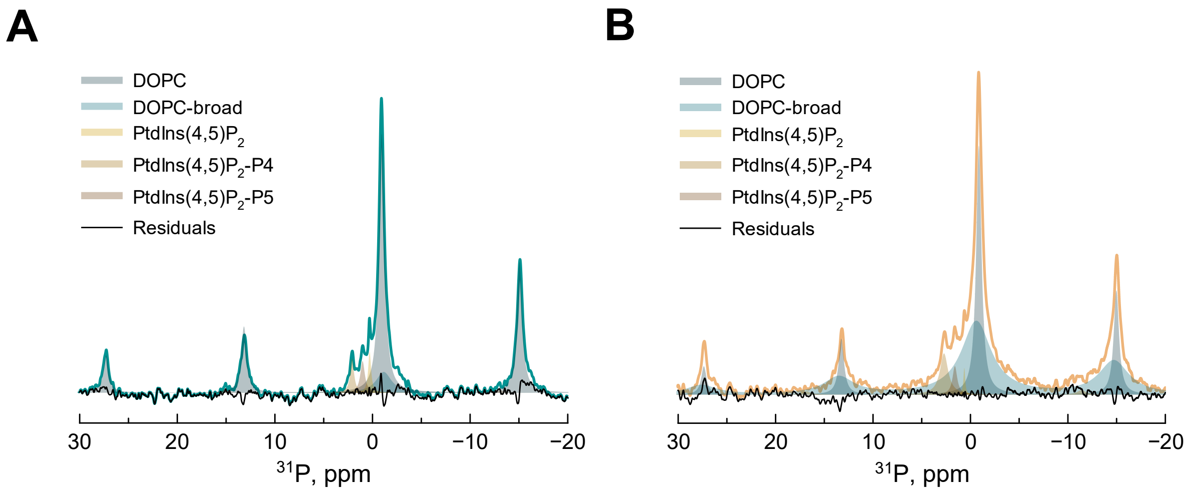


**Figure S9.** $^{31}$P direct excitation ssNMR spectral deconvolution of DOPC/PtdIns(4,5)$P_2$ liposomes in the absence (**A**) and presence of $CALM_{wt}$ protein (**B**). Spectra deconvolution was used to determine the CSA parameters, shown in ***Table S8***. The spectra were acquired at 290 K using a MAS speed of 4 kHz.

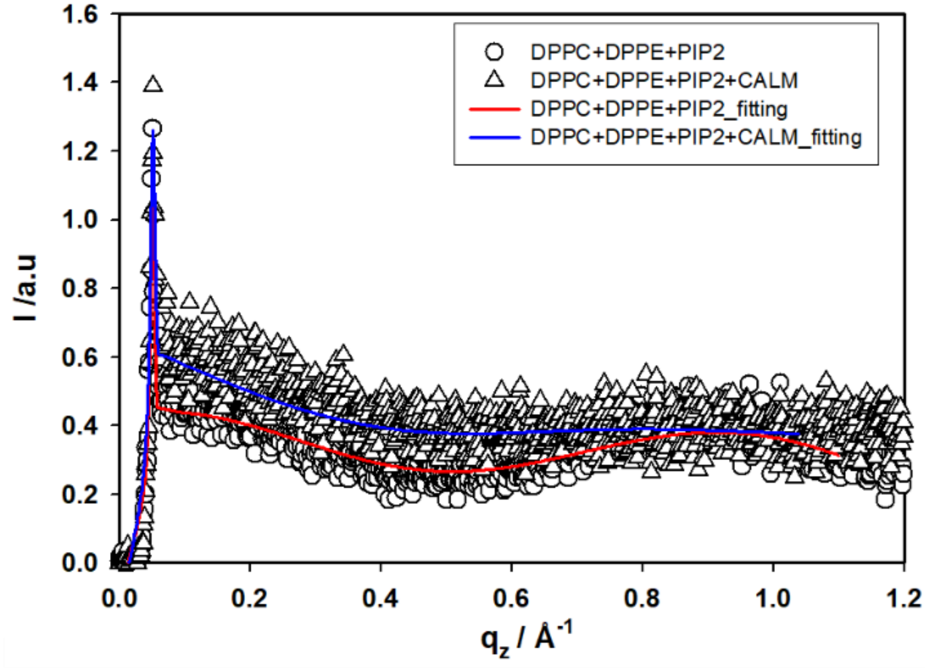


**Figure S*10*.** Bragg rod analysis.

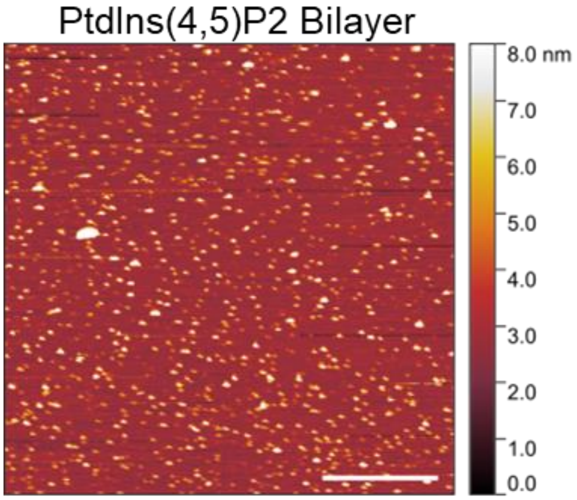


**Figure *S11.*** PtdIns4,5$P_2$ clusterization: SLBs composed by DOPC:DOPE:PtdIns4,5P2 in HKM buffer are examined by fluid-phase peak force tapping mode AFM. Scale bar is 0.5 mm.
